# The long noncoding RNA *FOSIL* shapes the dynamics of *FOS* induction

**DOI:** 10.64898/2026.11.04.764297

**Authors:** Juan Pablo Unfried, Igor Ulitsky

## Abstract

Immediate-early genes require rapid, transient, and precisely tuned transcriptional activation in response to extracellular signals, yet how this kinetic precision is achieved remains poorly understood. We characterize *FOSIL* (*LINC01220*), an inducible lncRNA locus downstream of *FOS*. Depleting *FOSIL* RNA delays heregulin (HRG)-induced *FOS* mRNA and protein accumulation and prolongs the response, indicating that *FOSIL* sharpens both the onset and the shutoff of the induction wave. Capture Micro-C maps show that *FOSIL* depletion weakens the chromatin boundary between *FOS* and *FOSIL* and increases local compaction across the pre-assembled *FOS*-*FOSIL* neighborhood during serum starvation, demonstrating that basal *FOSIL* production maintains a locally insulated, poised conformation prior to signal arrival. Upon HRG stimulation, *FOSIL* depletion blunts the rapid movement of the locus toward nuclear speckles, diminishes early speckle-proximal *FOS* transcription, and extends late firing at speckle-distant alleles. At single transcription active sites, nascent *FOSIL* activity correlates with closer speckle proximity and higher *FOS* output, and activating the endogenous *FOSIL* promoter by CRISPRa, but not supplying *FOSIL* RNA *in trans*, enhances both speckle association and *FOS* induction. Together, these findings support a mechanism where endogenous *FOSIL* activity coordinates pre-stimulus chromatin folding with dynamic nuclear speckle positioning to fine-tune the onset, amplitude, and resolution of immediate-early gene expression.

## Introduction

Immediate-early genes (IEGs), including *FOS*, respond to growth factor and neuronal inputs and can initiate subsequent gene-regulatory programs on very short timescales ^1,2^. MCF7 breast cancer cells respond to two ErbB ligands, with EGF promoting proliferation and transient ERK activation and heregulin (HRG) promoting differentiation and sustained ERK signaling ^3^. Despite these differences, both ligands induce a transient *FOS* mRNA pulse ^4,5^. How such induction waves are initiated, amplified and attenuated with precise and robust dynamics is a central question in transcriptional regulation.

Binding of EGF or HRG to cell-surface ErbB receptors activates the cytoplasmic RAS-RAF-MEK-ERK cascade, from which phosphorylated ERK and RSK2 translocate to the nucleus ^6,7^. In the nucleus, ERK activates ELK1, which acts with SRF at the *FOS* serum response element (SRE), while RSK2-dependent CREB phosphorylation provides a second *FOS* promoter input ^6–8^. Nuclear ERK activation is nevertheless transient after HRG, in part because DUSPs dephosphorylate the kinase ^4,6^. Depleting these phosphatases extends nuclear signaling but scarcely changes the duration of the mRNA pulse; blocking protein synthesis instead prolongs it ^4,7^. Under HRG, sustained cytoplasmic ERK activity stabilizes the *FOS* protein, allowing its accumulation to outlast the RNA response ^4,5^.

Candidate regulatory elements at the human FOS locus have been predicted ^9^ but none has been systematically tested for its effect on human *FOS* expression. Noncoding transcription can contribute to local regulation and is proposed to be a prominent early component of stimulus-responsive programs ^10,11^. In neurons, enhancer RNAs facilitate release of paused polymerase at immediate-early genes ^12,13^. Likewise, long noncoding RNA (lncRNA) loci have been shown to shape stimulus-responsive transcription of nearby coding genes ^14,15^. Local regulation by lncRNA loci can involve promoter DNA, transcription or splicing, often without a specific role for the mature RNA product ^16–18^. Where RNA itself is involved, local concentration and amplification can explain strong effects despite low transcript abundance ^19^. Induction of a noncoding locus could therefore change the local regulatory environment in which a nearby gene responds to stimuli, but the direct mechanistic consequences of this transcription remain unclear.

At the chromatin conformation level, spatial interaction contacts are not necessarily correlated with transcription. Many enhancer-promoter contacts in mouse embryonic stem cells persist after acute transcriptional inhibition, while disrupting higher-order contacts in differentiating human cells has comparatively modest effects on gene activation ^20,21^. During neuronal activation, however, recent higher-resolution contact maps reveal rapid formation of chromatin loops at immediate-early genes where enhancer RNAs have been implicated as key functional drivers of the local architectural rewiring ^22^. However, it remains unclear how inducible noncoding loci influence the organization of their neighbors before stimulation, in the lead-up to transcription activation and as transcription proceeds.

Gene regulation can also depend on the locus position relative to nuclear speckles ^23,24^. These membraneless compartments concentrate RNA-processing factors in their periphery, including the U1 and U2 snRNAs ^25^. The speckle periphery is also enriched in the lncRNA *MALAT1*, which can act as a biophysical catalyst, enhancing the recruitment of SRSF1 to elongating Pol2 during hypoxia-induced alternative splicing ^26^. At the inner core of speckles, SON and SRRM2 organize and are essential for speckle integrity ^25,27^. Active genomic regions have been found to associate with speckle-near compartments, where proximity can enhance RNA splicing and the expression of selected target genes ^23,28,29^. Transcription factors can promote speckle association through both transcription-dependent and transcription-independent activities ^30^. Interestingly, perturbing chromatin architectural factors CTCF or cohesin can also alter speckle association ^31,32^, and CTCF clustering around speckles can lead to architectural changes that direct alternative splicing choices during senescence ^33^. Thus, transcriptional and co-transcriptional activities, and nuclear positioning may influence one another; yet it is unclear how their integration eventually determines a specific regulatory program and whether and how an inducible noncoding locus can contribute to this relationship during or in preparation for rapid transcriptional response.

Here, we investigate *FOSIL* (*LINC01220*), a previously uncharacterized lncRNA locus found 15.6 kb downstream of *FOS* and induced upon growth factor stimulation. We first define the 3D neighborhood and then combine depletion, ectopic expression and endogenous activation with expression, Capture Micro-C and RNA and DNA imaging to identify how *FOSIL* shapes the timing and magnitude of *FOS* induction. Depleting *FOSIL* RNA delays the bulk induction wave, and leads to increased local chromatin compaction before stimulation and reduced speckle proximity after HRG. Ectopic expression does not enhance induction, pointing to the nascent RNA or its production as the functional elements, whereas endogenous activation increases *FOS* output and speckle-proximal association of active loci. We propose that *FOSIL* transcription specifies the degree of local chromatin compaction needed to support a coordinated and tight transcription pulse and RNA processing near nuclear speckles, enabling timely FOS protein induction.

## Results

### *FOSIL* is a distinct, inducible transcriptional unit within the *FOS* regulatory landscape

*FOS* induction has been studied for four decades ^1,34,35^ with studies mostly focusing on the DNA elements in the locus, FOS promoter and several upstream enhancer elements in a mostly gene-sparse locus. In humans, the ∼250 kb genomic region surrounding *FOS* contains only one other spliced gene model, transcribed from a promoter 15.5 kb downstream of FOS and on the same strand, with no known role, that we have termed *FOSIL* (<u>FOS</u>-related Intergenic <u>L</u>ncRNA, annotated as *LINC01220*, Figs. 1a and S1a).

**Figure 1.**
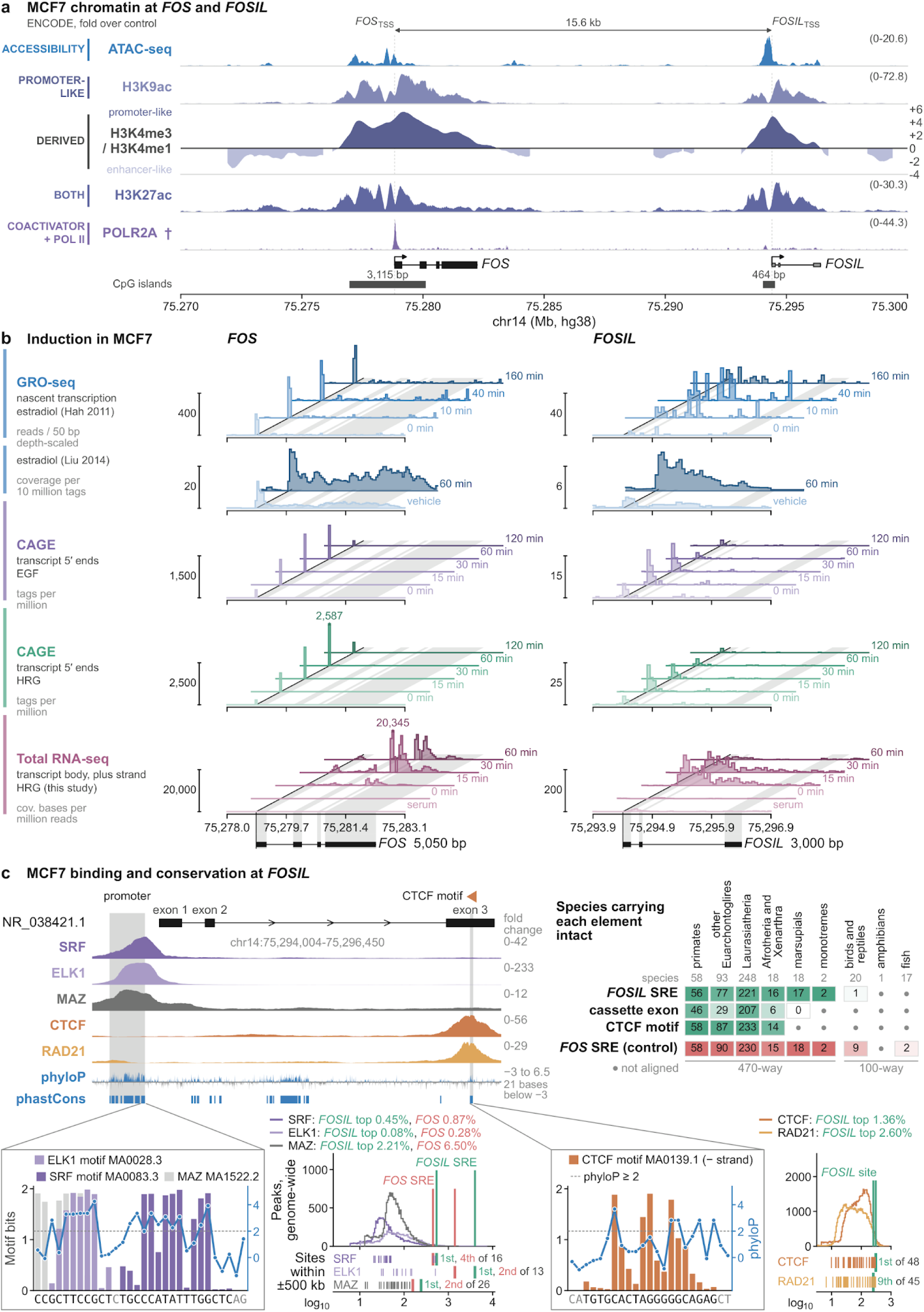
*FOSIL* is a distinct, inducible transcriptional unit within the *FOS* regulatory landscape. **(a)** MCF7 chromatin accessibility, histone marks and POLR2A occupancy across *FOS*-*FOSIL* (GRCh38), with RefSeq/MANE models, CpG islands and the log2 H3K4me3:H3K4me1 ratio. ENCODE tracks use independent fold-change-over-control scales (†, POLR2A growth conditions unspecified). **(b)** Plus-strand GRO-seq after estradiol (0-160 min, two pooled replicates, or 1 h versus vehicle, mean of three), FANTOM5 CAGE after EGF or HRG (three libraries per induced state, two at the shared baseline), and HRG total RNA-seq in CTRL GapmeR-treated cells (two libraries per state, beginning with normal serum [S] and serum starvation [0 min]). Carets mark clipped *FOS* peaks. **(c)** MCF7 SRF, ELK1, MAZ, CTCF and RAD21 ChIP-seq, vertebrate phyloP and phastCons elements across *FOSIL* (top left, chr14:75,294,004-75,296,450, NR_038421.1). Shading marks the promoter SRE (SRF CArG box with upstream ELK1 ETS site) and exon-3 CTCF motif (MA0139.1, minus strand). Top right, numbers give the count and shading the fraction of species in each clade with an intact SRE CArG box, both intron-2 cassette-exon splice sites or the exon-3 CTCF motif. The *FOS* SRE is shown for comparison. Mammals were scored in the 470-way alignment and other vertebrates in the 100-way alignment (dot, unaligned). Below, motif information content (bars) and vertebrate phyloP (dots, dashed line at 2) at the ELK1, SRF, MAZ and CTCF sites. Adjacent histograms and ticks place the *FOSIL* peaks (green) and *FOS* SRE peaks (coral) among MCF7 peaks genome-wide and within 500 kb, respectively. See also Figure S1.

Public transcriptomic and epigenomic data in MCF7 cells show a distinct accessible, acetylated start region with H3K4me3 enrichment relative to H3K4me1, as expected for an independent promoter, and detectable POLR2A (Fig. 1a). In FANTOM5 CAGE data, *FOSIL* responds to treatment with the growth factors EGF and HRG with an approximately threefold increase in capped RNA signal at 15 min, with the maximum occurring earlier than for *FOS*. GRO-seq after estradiol showed an approximately 11-fold increase in nascent transcription at 40 min, in two independent datasets ^36,37^. No readthrough evidence connects the two genes, supporting the model that FOS and FOSIL are two independent transcriptional units (Fig. S1b). Additionally, our RNA-seq data support the formation of two splice junctions of the curated three-exon model (NR_038421.1), indicating that the locus is spliced but expressed at low absolute level in unstimulated cells. Annotated alternative cassette-exon and distal fourth-exon junctions were undetectable by RNA-seq.

We also examined the broader ∼250 kb region, and identified regions that correspond to those studied in the mouse *c-fos* enhancer studies ^38,39^ that we will denote correspondingly e1-e5; e5 includes the *FOSIL* start site (Fig. S1a) ^39^. In our total-RNA sequencing time course of HRG-treated cells, mean normalized counts increased 45.5-fold across the 2,007-bp NR_038421.1 genomic span and 18.5-fold across its 490-bp exon union in CTRL cells between 0 and 15 min after HRG treatment, and *FOSIL* and *FOS* reached their sampled maxima at 15 and 30 min, respectively (Fig. 1b; two biological replicates). No other lncRNA production was noted throughout the locus. FOSIL thus corresponds to a growth factor-inducible and partially spliced lncRNA co-induced with FOS, while being transcribed at overall lower levels.

### Focal conserved elements overlap the *FOSIL* locus

We next examined conservation of the locus. An annotation-based survey of 15 vertebrates placed *TMED10*, *FOS* and *JDP2* on one chromosome in every examined species of vertebrates through bony fish, but identified a same-strand noncoding candidate between *FOS* and *JDP2* only in eutherian mammals, including all six surveyed species with RNA-seq data (human, mouse, rat, cat, dog, ferret) (Fig. S1c). Mean phyloP across the human 2,587-bp analysis interval was not significantly higher than length-matched local intervals at four alignment depths, suggesting overall poor sequence conservation, in line with most lncRNA genes found away from protein-coding genes ^40^. Overall, *FOSIL* conservation is ordinary for the lncRNA class (ranked at the 53.7th percentile of 18,534 GENCODE lncRNA exon unions) (Fig. S1d) and the 40.5th percentile among other genes with three detected sequence orthologs (Fig. S1f). Despite poor sequence conservation, several conserved elements detected by phastCons overlap the locus, covering 6.7% of exonic and 12.4% of non-exonic bases (Fig. 1c). These include a 103-bp intron-2 element, covering most of an annotated 69-bp cassette exon that is absent from NR_038421.1 and not transcribed at any appreciable level in MCF7 cells (about 1% of *FOSIL* exon-2 splicing in public tissue and tumor RNA-seq), and an element in exon 3 overlapping CTCF and RAD21 ChIP-seq peaks (Figs. 1c and S1a). Underlying this peak is a consensus CTCF binding motif, with 16 of its 19 bases showing evolutionary constraint. This motif is highly occupied by CTCF in MCF7 cells, ranking in the top 1.4% genome-wide across eight ENCODE experiments, and it is the strongest CTCF peak in a 1Mb window. In the same region, the RAD21 peak ranks in the top 2.2-2.6% in both available experiments (Fig. 1c). At the *FOSIL* promoter, a constrained element contains a CArG box 85 bp upstream of the *FOSIL* TSS, which is bound by SRF and ELK in MCF7 ChIP-seq data. With an ETS site just upstream, this CArG box has the layout of the *FOS* serum response element (SRE) ^41,42^. These SRF and ELK1 peaks rank in the top 0.45% and 0.08% genome-wide and are the strongest within 1Mb, with stronger evidence of binding than the peaks at the *FOS* promoter SRE (top 0.87% and 0.28%) (Fig. 1c).

In mouse and rat, the aligned sequence of *FOSIL* exon 3 corresponds to the last exon of annotated lncRNAs *Gm60138* and *LOC148005395*, respectively. In mouse, the CTCF motif is intact and occupied in every ENCODE CTCF experiment, and in adult liver it is bound by CTCF in human, macaque, mouse, rat and dog ^43^.

The CArG box of the *FOSIL* SRE is intact in every mammal group, monotremes included, as is that of the *FOS* SRE. The CTCF motif and the cassette exon’s splice sites are intact in placental mammals; beyond them the motif is not aligned in any species (Fig. 1c). The DNA elements in the FOSIL region are thus highly constrained, while the rest of the RNA locus is evolving rapidly, but appears to maintain lncRNA production in mammals.

### Endogenous *FOSIL* activity shapes the timing and magnitude of *FOS* induction

To test the role of *FOSIL* during growth factor treatment, we decided to focus on HRG, as it induces both *FOS* and *FOSIL* the strongest and also does not affect proliferation ^4,5^, which could become a confounding factor for expression measurements at later time points. We measured FOSIL and FOS levels by RT-qPCR, RNA-sequencing and flow cytometry after treatment with GapmeR ASOs designed to degrade FOSIL or control GapmeRs. We profiled the normal fed state (S), serum-starved (0) and in a time chase after HRG treatment at 15, 30, 60, 90, and 180 minutes (Fig. 2a,b). We used L1 and L2 LNA-GapmeRs, which direct RNase H cleavage and therefore act on nascent as well as mature RNA ^46^, targeted at distinct sites in *FOSIL* RNA and reduced its baseline level by 55% or higher, and kept it lower after HRG (Fig. 2c-f) relative to a control non-targeting GapmeR (CTRL). Each GapmeR matched the transcriptome only at its *FOSIL* site, and no gene carried a near match for both, so changes shared by L1 and L2 are unlikely to arise from a common off-target (Fig. S2a-c; Methods). In CTRL cells, total *FOSIL* peaked 15 min after HRG at about 40 times its serum level and then declined, whereas fully spliced *FOSIL* rose less, to about 16 times, and peaked later, at 30-60 min, consistent with the CAGE data (Fig. 1b), and with overall fast induction dynamics followed by co-transcriptional processing (Fig. 2c,e). Transcriptome-wide, very few genes per time point differed significantly from CTRL by more than 2-fold with either GapmeR (Fig. S2d); none did when each GapmeR’s change from its own serum sample was compared with CTRL’s, supporting their specificity to *FOSIL* (Methods).

**Figure 2.**
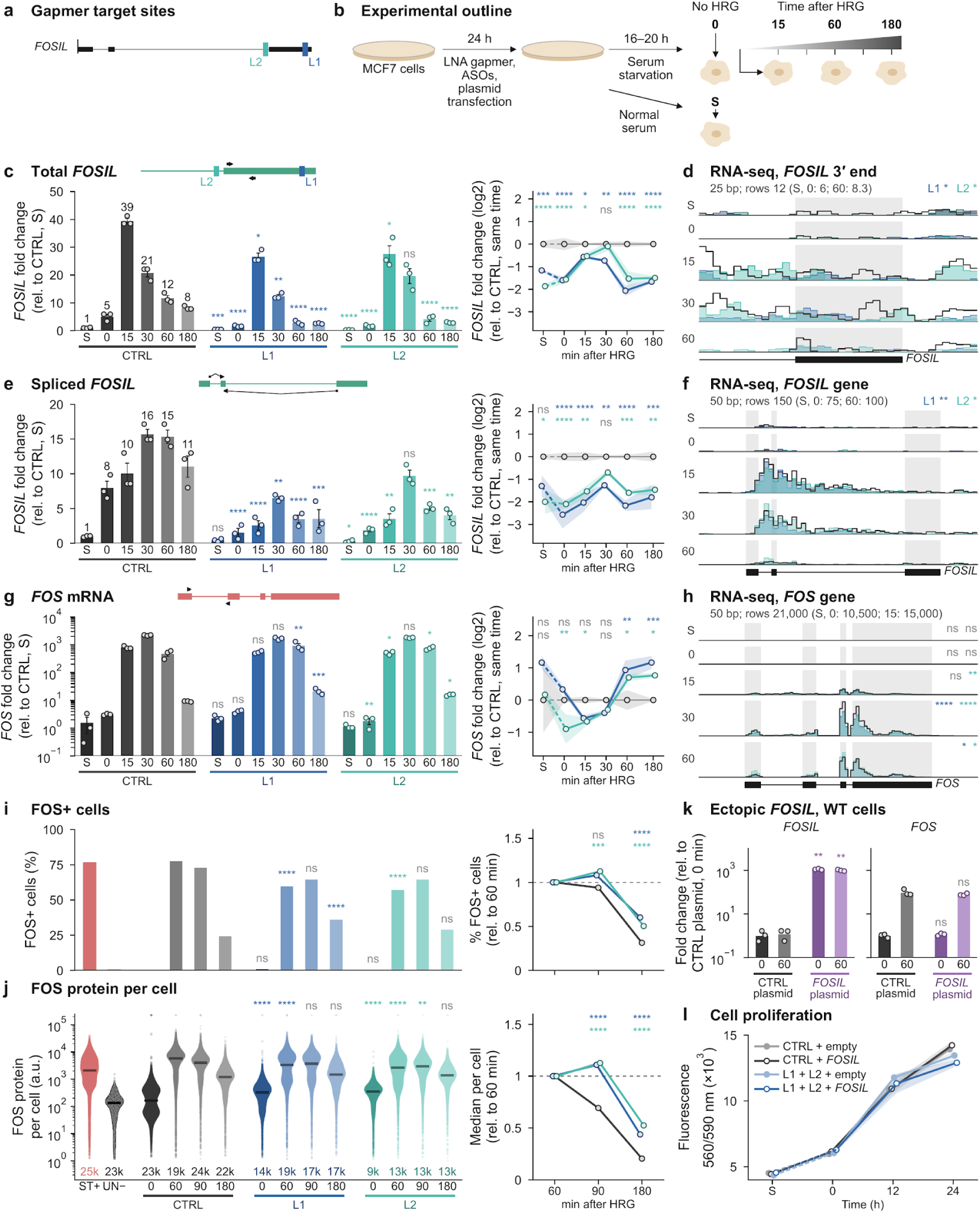
Endogenous *FOSIL* perturbation alters the *FOS* induction wave. **(a)** LNA-GapmeR target sites (L1 and L2) on *FOSIL*. **(b)** Experimental timeline in normal serum (S), after serum starvation (0 min) and 15-180 min after HRG. **(c,e,g)** RT-qPCR of total *FOSIL* (c), spliced *FOSIL* (e) and *FOS* mRNA (g) after CTRL, L1 or L2 GapmeR treatment, normalized to *ACTB*. Left, expression relative to CTRL in serum (mean ± SEM, circles show three biological replicates). Right, log2 fold change relative to time-matched CTRL (bands, ± SEM for each condition). Primer positions are shown above. **(d,f,h)** Total RNA-seq coverage across the *FOSIL* 3′ end (d), *FOSIL* locus (f) and *FOS* locus (h) from S to 60 min (two biological replicates pooled per condition and state). Bin sizes and row scales are indicated above. **(i,j)** Flow cytometry of FOS protein in CTRL, L1 and L2 cells before HRG and at 60, 90 and 180 min after HRG (one pooled sample from three independent transfections per condition and time point, cell counts in thousands). **(i)** FOS-positive cells (%). ST+ and UN− mark stained and unstained 60-min HRG controls from a separate run. **(j)** Single-cell FOS fluorescence intensity (bars, medians). Insets show values relative to each condition’s 60-min level. **(k)** RT-qPCR of *FOSIL* and *FOS* at 0 and 60 min in WT cells transfected with ectopic *FOSIL* or empty vector, normalized to *GAPDH* and relative to the 0-min vector control (geometric means, n = 3). **(l)** PrestoBlue fluorescence after cotransfection with CTRL or pooled L1 + L2 GapmeRs and empty or *FOSIL* plasmid, in serum, after starvation and at 12 and 24 h after HRG (mean ± SEM, four wells across two independent experiments). In c,e,g, two-way ANOVA on ΔCt was followed by Dunnett comparisons with time-matched CTRL, with S tested separately. RNA-seq comparisons used Gart’s one-sided exact test (mid-P, five time points pooled) in d,f and Benjamini-Hochberg-adjusted DESeq2 Wald tests in h. In i,j, one-sided minimum-effect tests (|h| > 0.2 in i, |d| > 0.2 in j) compared each GapmeR with CTRL at each time and for the change from 60 min (insets), with Holm correction. In k, Welch’s t-tests on ΔCt were Holm-adjusted. *P < 0.05, **P < 0.01, ***P < 0.001, ****P < 0.0001. ns, not significant. See also Figure S2.

In FOSIL-depleted cells, FOS was still induced by HRG to levels similar to those in control cells, but with altered dynamics, which included a delay in the induction and higher levels at the later time points (Fig. 2g,h). Mean *FOS* abundance was lower than CTRL with either GapmeR at 15 and 30 min and 1.6 to 2.2 times higher at 60 and 180 min (Fig. 2g), and was likewise lower by RNA-seq at 15 and 30 min and trending higher by 60 min (Fig. 2h). In a separate experiment, FOS protein measured by flow cytometry peaked later and lower in cells treated with either FOSIL-targeting GapmeRs and persisted longer than in control-treated cells (Fig. 2i,j). These depletion experiments show that intact *FOSIL* transcription or its RNA product regulate the timing and the persistence of FOS induction.

To test the potential contribution of ectopically provided FOSIL RNA, we transfected the cells with a FOSIL-expressing plasmid, which led to supraphysiological *FOSIL* RNA levels at 859-to 1,128-fold the level of time-matched vector controls, yet this did not affect *FOS* induction (Fig. 2k). Because the RNA was supplied *in trans*, this points to the functional element in the locus being RNA production or processing rather than the amount of RNA present, under the tested conditions. Proliferation over 24 h was similar with or without the GapmeRs or ectopic *FOSIL,* showing that HRG-induced FOS, even when altered by FOSIL-targeting, does not affect proliferation (Fig. 2l).

### Capture Micro-C maps define the *FOS*-*FOSIL* contact neighborhood

To establish the chromatin context in which *FOSIL* acts to regulate *FOS* induction, we mapped contacts across the ∼340 kb *FOS*-*FOSIL* extended neighborhood by Capture Micro-C, which profiles DNA fragments spatially proximal *in situ* in MCF7 cells treated with the non-targeting control GapmeR (CTRL). We profiled four biological replicates per state: normal serum (S), serum-starved just before HRG (0), and 15 and 60 min after HRG. Micro-C revealed a prominent insulated domain of ∼175 kb formed between two flanking CTCF sites left and right (CTCF_L_ and CTCF_R_), found 146.2 kb upstream of FOS and another 13.6 kb downstream of FOSIL, respectively (Fig. 3a). We annotated twelve contacts supported by significantly more read pairs than expected at their genomic separation, using × between the names of the paired landmarks. These were detected in every state and we numbered them 1-12 by their support strength in the normal serum (Fig. 3a,b). We also annotated five boundaries, regions that have few contacts across them, which correspond to dips in an insulation score measuring contacts crossing each position (Fig. 3c). To refer to boundaries we use a vertical bar (|) preceding a single landmark or separating two landmarks. The five boundaries are |e2 core, e4|*FOS*, *FOS*|*FOSIL*, |*FOSIL*_TSS_ and |CTCF_R_ (Fig. 3d,f). We also refer to two regions as “intervals”, defined as regions between adjacent boundaries ^47,48^, the 6 kb *FOS* gene interval (→*FOS*) and the 11 kb interval between *FOS* and the *FOSIL* promoter (*FOS*→*FOSIL*; Fig. 3f,g). To test for changes between states, libraries were normalized by depth and composition using TMM ^49^ and changes in read counts were evaluated with edgeR, as used by the diffHic method ^50,51^, with changes called as significant if they had a <1% false discovery rate (Methods).

**Figure 3.**
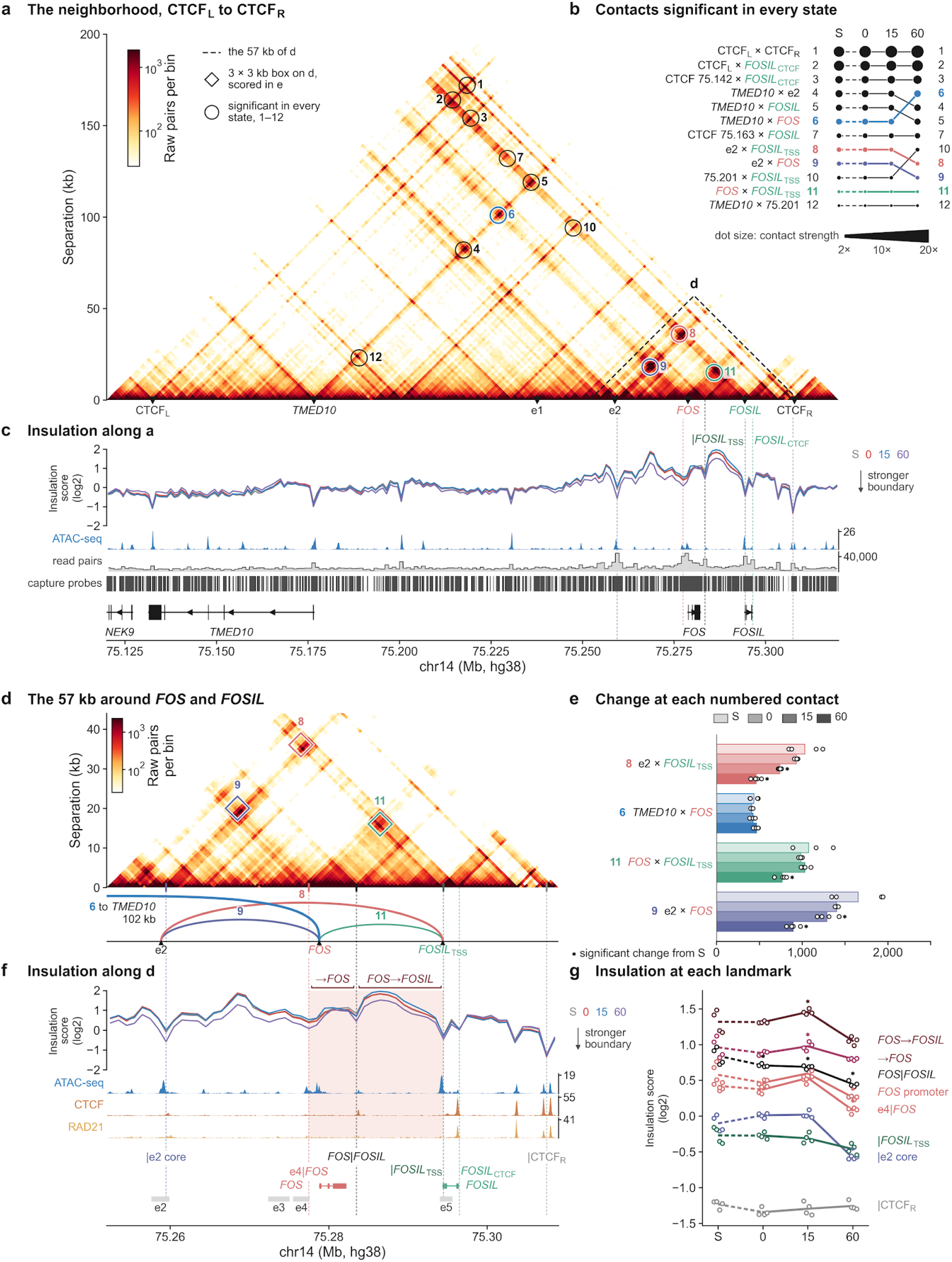
The contact neighborhood of *FOS* and *FOSIL*. Capture Micro-C in CTRL GapmeR-treated MCF7 cells in normal serum (S), after starvation (0) and at 15 and 60 min after HRG (four biological replicates per state). **(a)** Pooled contact map across the *FOS*-*FOSIL* neighborhood from CTCF_L_ to CTCF_R_ (16 libraries, 1-kb resolution). Circles mark the 12 contacts detected in all four states, numbered by serum contact strength (observed over distance-expected counts). The dashed box marks the region in d. **(b)** Contact ranks across states. Circle size indicates strength, and numbers at right identify contacts in rank order at 60 min. **(c)** Insulation profiles across a. Dashed vertical lines mark five boundaries (|e2 core, e4|*FOS*, *FOS*|*FOSIL*, |*FOSIL*_TSS_ and |CTCF_R_) and the exon-3 CTCF site (*FOSIL*_CTCF_), above MCF7 ATAC-seq, read-pair coverage and capture-probe positions. **(d)** Pooled map of the 57-kb region around *FOS* and *FOSIL* at 500-bp resolution. Diamonds mark 3 × 3-kb contact boxes, for contacts 8, 9 and 11, with corresponding arcs below. **(e)** TMM-normalized read pairs per million in-window pairs for contacts 8 (e2 × *FOSIL*_TSS_), 6 (*TMED10* × *FOS*), 11 (*FOS* × *FOSIL*_TSS_) and 9 (e2 × *FOS*). **(f)** Insulation profiles across d, with MCF7 ATAC-seq, CTCF and RAD21 ChIP-seq, elements e2-e5 (grey bars), and the →*FOS* and *FOS*→*FOSIL* intervals (shaded). **(g)** Insulation at the five boundaries, the *FOS* promoter and the two intervals (mean over interior bins). Lower scores indicate stronger insulation. In e,g, open circles show individual libraries and bars (e) or lines (g) show means. Asterisks mark changes from serum (diffHic/edgeR quasi-likelihood F-tests, FDR < 1%). Interval tests count pairs with both ends inside. See also Figure S3.

The two strongest contacts join CTCF_L_ to CTCF_R_ (15.8-fold enrichment) and to the *FOSIL* exon-3 CTCF site (15.2-fold; Fig. 1c), and nine of the 12 have one end at either *FOS* or *FOSIL* (Fig. 3b, S3d). In serum, the *FOS* promoter’s top contacts include e2 (an element with prominent enhancer marks and enhancer RNA transcription ^45^, Fig. S1a), the *TMED10* promoter (upstream neighboring gene) and the *FOSIL*_TSS_ (Fig. 3a,b). The seven strongest of ten named contacts were also the strongest in an independent publicly available genome-wide WT MCF7 Micro-C map, supporting a robust chromatin conformation formed in fed cells ^52^ (Fig. S3a-c). Starvation and HRG treatment did not raise contacts uniformly; instead, they redistributed the contacts selectively among e2 and the two promoters (Fig. 3e). The e2 × *FOS* and e2 × *FOSIL*_TSS_ contacts were weakened after HRG, with each falling to about half its serum read pairs by 60 min. There was also a decrease in the *FOS* × *FOSIL*_TSS_ contact at 60 min. The *FOS* promoter also gained contact with CTCF_L_ and lost contact with e3 by 60 min. The early induction wave therefore occurred within this pre-formed contact neighborhood, without a general increase in contacts between the *FOS* promoter and its neighbors, and with specific contacts weakening over the induction time.

When considering boundaries, we also observed a combination of changing and fixed landmarks (Fig. 3f,g). During starvation and the first 15 min, the *FOS*|*FOSIL* boundary strengthened, and remained stronger through 60 min, while other boundaries remained steady. Contact frequencies within both intervals rose between 0 and 15 min, coinciding with the early *FOSIL* RNA induction (Fig. 1b), and the gain was gone by 60 min. By then insulation was strongest among all examined timepoints for most boundaries.

Together, the maps show that *FOS* is induced within a *FOS*-*FOSIL* neighborhood already assembled in a normal fed state accompanied by selective redistribution of local contacts and anchored at CTCF sites, notably without formation of new contacts or boundaries. Conformation changes in the region begin during starvation, before HRG stimulation (Fig. 3g).

### *FOSIL* RNA restrains local compaction and contact enrichment across the *FOS* neighborhood from starvation and early after HRG induction

We then tested how the chromatin conformation changes after FOSIL depletion, using the same Micro-C capture set at S, 0, 15 and 60 min (Fig. 4). Depleting *FOSIL* RNA increased contact inside this pre-assembled neighborhood, mostly in the starved state before HRG, while its outer boundary exhibited limited changes. Of the 37 *FOS*-promoter contacts and 44 insulation positions that changed significantly, most fell in groups whose change appears by 0 min, before HRG treatment, and then holds or declines; only two contacts peaked transiently at 15 min around e1, and five rose gradually to 60 min downstream of *FOSIL* (Fig. 4a-b,f). We also measured the two intervals, the *FOS* gene and the stretch to the *FOSIL* promoter (Fig. 3f,g). A higher mean insulation score over an interval, which indicates enriched contacts across it, was taken as greater local compaction, following Micro-C studies linking contact enrichment with chromatin compaction across yeast, mouse and human cells ^53–57^. The named contacts of the locally constrained exon-3 CTCF site were followed in each state (Fig. S4).

**Figure 4.**
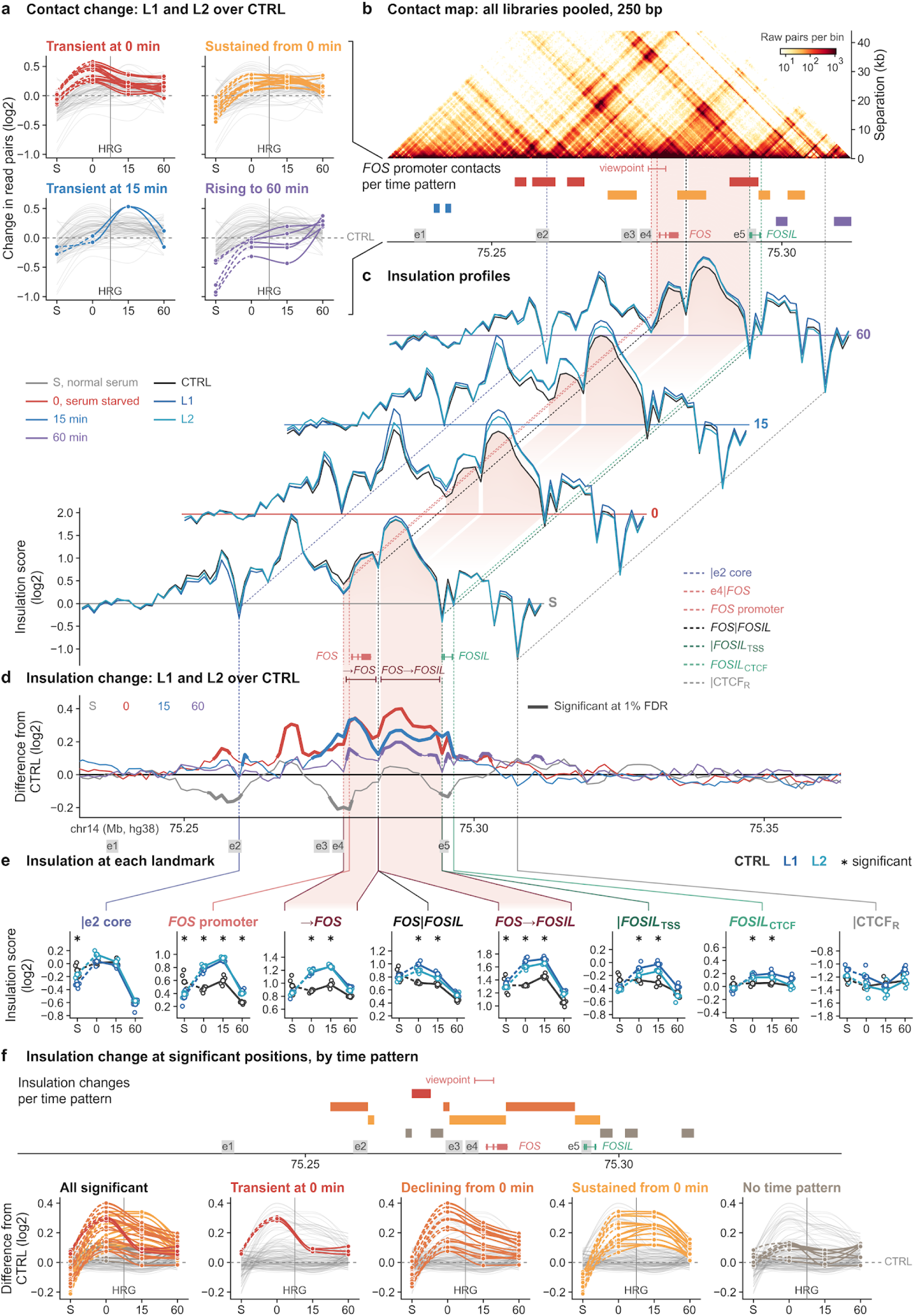
*FOSIL* sets local chromatin compaction from *FOS* to the *FOSIL* promoter in starved cells and during early *FOS* induction. **(a)** Log2 fold change in read pairs at *FOS*-promoter contacts for L1 and L2 together versus time-matched CTRL, in normal serum (S), after starvation (0) and at 15 and 60 min after HRG. Contacts that changed significantly are grouped by time course into transient at 0 min, sustained from 0 min, transient at 15 min or rising to 60 min. **(b)** Capture Micro-C map pooled across all 48 CTRL, L1 and L2 libraries (250-bp bins, chr14:75,232,000-75,312,000). Below the map, colored bars mark the *FOS*-promoter contact positions in a, above elements e1-e5 and gene models. **(c)** Mean insulation profiles for CTRL, L1 and L2 (four biological replicates per condition and state). Shading marks the →*FOS* and *FOS*→*FOSIL* intervals. **(d)** Insulation difference at each time point (mean of L1 and L2 minus CTRL). Thick segments mark significantly changed bins. **(e)** Insulation at six landmarks and the two intervals. Open circles show individual libraries and lines show means. Asterisks mark time points at which L1 and L2, tested together, differed from CTRL. Higher scores indicate weaker insulation. **(f)** Insulation change at significant positions, shown together and grouped by time course into transient at 0 min, declining from 0 min, sustained from 0 min or without a detectable time pattern. Colored bars above the plots mark genomic positions. In a and f, thin grey lines show the other positions and the dashed zero line marks CTRL. Changes were tested with diffHic/edgeR quasi-likelihood F-tests (FDR < 1%); temporal profiles were grouped by k-means clustering (Methods). See also Figure S4.

In control cells both intervals become transiently more compact at 15 min (Fig. 3g). In serum, the stretch from the boundary downstream of *FOS* to the *FOSIL* promoter already had 8.1% more read pairs in GapmeR-treated cells than in CTRL, while insulation at the *FOS* promoter, “|e2 core” and “|*FOSIL*_TSS_” was stronger. From starvation onward, every GapmeR-treated library scored above every CTRL library on both intervals. The largest changes were concentrated in these two regions, with smaller changes extending up to the e2 region and down to the *FOSIL* gene body (Fig. 4d,e). The *FOS* promoter’s contacts with the *FOSIL*_TSS_, e2 and the exon-3 CTCF site were furthest above CTRL at 0 min (Fig. 4a,b); both *FOSIL* anchors also gained contact with their other partners (Fig. S4). When considering boundaries, the *FOS*|*FOSIL* boundary, which strengthened with starvation in control cells, was weakened by FOSIL depletion at 0 and 15 min. At 15 min, 8.5% more read pairs crossed the exon-3 CTCF site itself, while the outer boundary, |CTCF_R_, showed no detected difference from CTRL in any sampled state (Fig. 4e). From starvation onward, intact FOSIL production restrains this compaction and the broader contact environment around *FOS*; without it, both intervals are more compact and contact enrichment increases between the two promoters and with several neighboring partners (Fig. 4a,b,e).

Without the RNA, both intervals are already more compact at 0 min, before HRG, preceding the delayed bulk FOS response (Figs. 2g,h and 4). We propose that, from starvation onward, *FOSIL* RNA production limits local folding of FOS to maintain an organization permissive for timely *FOS* activation.

### *FOSIL* perturbations alter *FOS* RNA output and reduce speckle proximity after induction

To study the dynamics of the response of FOS and FOSIL to growth factor at the level of individual active loci and with information about the spatial positioning of the loci, we next used a combination of RNA single molecule fluorescence in situ hybridization (RNA smFISH) and DNA FISH. In the first set of experiments, we used multiplex RNA HCR (the L series) to measure intronic *FOSIL* signal (preFOSIL), intronic *FOS* signal (preFOS) and *FOS* exon signal at detected transcription sites 15, 30 and 60 min after HRG (Fig. 5a,b). We focused on FOS transcription active sites (TAS) defined by a combination of the signals of preFOS and FOS exons. We observed five detected active sites at 15 and 30 min when transcription is active in most WT nuclei (representative images in Fig. 5b), consistent with five FOS DNA copies in MCF7 cells as detected by DNA FISH (see below). At 60 min some late-firing loci can still be detected but at much smaller numbers which hinders statistical analysis in most cases (Fig. 5b). *FOS* and *FOSIL* expression is very low before induction in the fed state (S) and serum-starved cells (0 min) and at 180 min after HRG (Fig. S5a), with no detectable active sites to identify the locus by RNA-HCR which confirms the quantifications in Fig. 2 and provide validation to the specificity of the probes. We therefore restricted active-site measurements mainly to 15 and 30 min, and analyzed also 60 min where possible.

**Figure 5.**
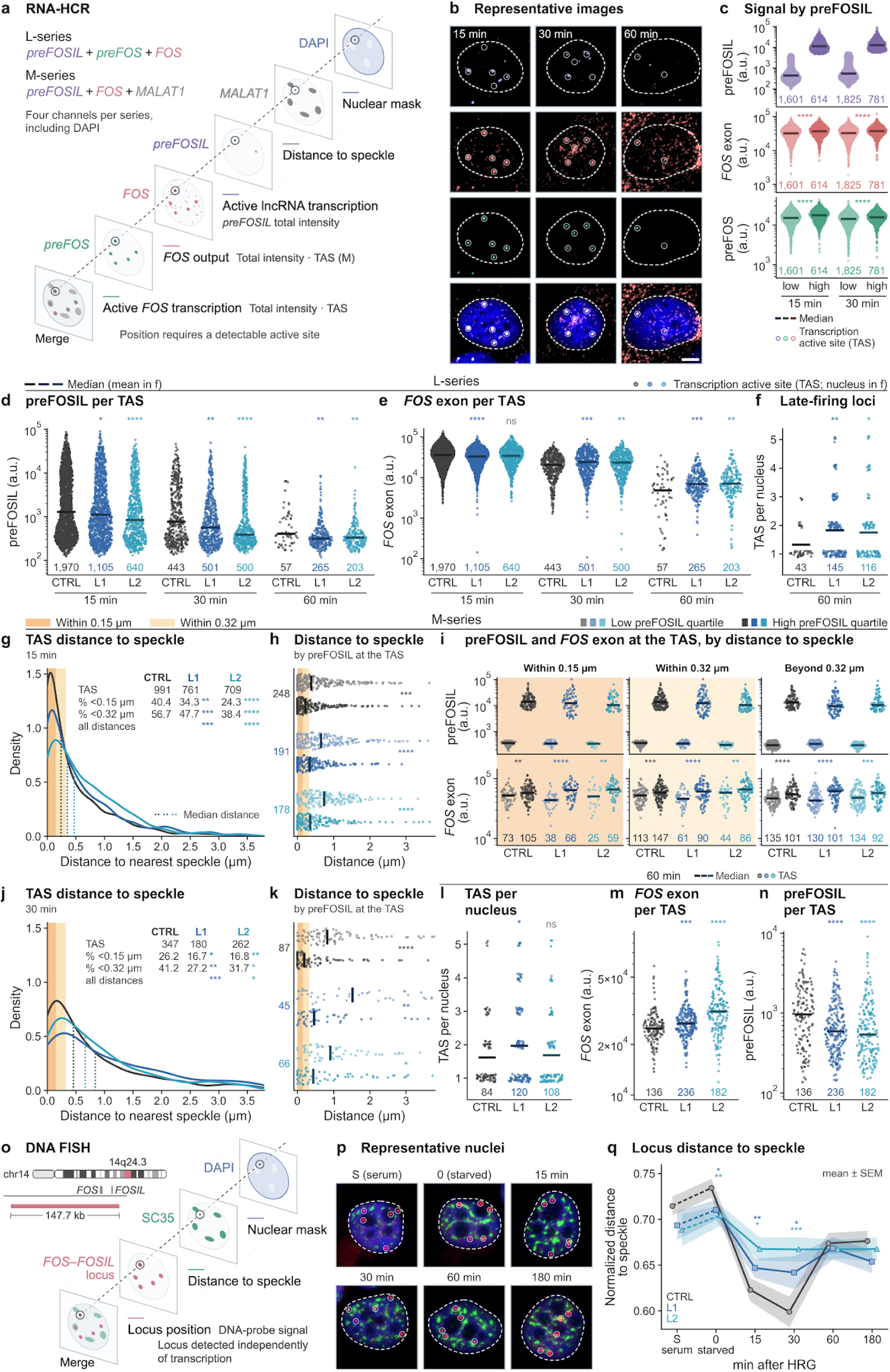
*FOSIL* perturbations alter RNA output and reduce early speckle proximity. **(a)** Multiplex RNA-HCR probe sets imaged with DAPI for the L series (preFOSIL, preFOS and *FOS* exon) and M series (preFOSIL, *FOS* exon and *MALAT1*). **(b)** Representative WT nuclei from the L series, with individual channels and merged images at 15, 30 and 60 min after HRG. Circles mark transcription active sites (TAS), dashed lines mark nuclear outlines, and the scale bar is 5 µm. **(c)** preFOS and *FOS* exon intensity per TAS in WT cells. Sites were split at the mean preFOSIL intensity at 15 and 30 min; the top row shows the split. **(d,e)** preFOSIL (d) and *FOS* exon (e) intensity per TAS in CTRL, L1 and L2 GapmeR-treated cells at 15, 30 and 60 min (L series). **(f)** Number of active *FOS* loci per active nucleus (at least one TAS) at 60 min. **(g)** Two-dimensional distance from each TAS to the nearest *MALAT1*-defined speckle at 15 min (M series). Dotted lines mark medians; the table gives TAS counts and percentages within 0.15 and 0.32 µm (shaded). **(h)** TAS-to-speckle distance at 15 min for the bottom (light) and top (dark) preFOSIL intensity quarters within each condition. **(i)** preFOSIL and *FOS* exon intensity in these quarters by speckle distance (within 0.15 µm, within 0.32 µm cumulative, or beyond 0.32 µm). **(j,k)** As in g,h, at 30 min. **(l-n)** Active *FOS* loci per active nucleus (l), *FOS* exon intensity per TAS (m) and preFOSIL intensity per TAS (n) at 60 min (M series). **(o)** DNA immuno-FISH with a 14q24.3 probe spanning *FOS* and *FOSIL*, SC35 immunostaining and DAPI. **(p)** Representative CTRL nuclei (red, *FOS*-*FOSIL* locus; green, SC35; blue, DAPI). **(q)** Distance from each DNA-FISH locus to the nearest SC35-marked speckle, normalized to the mean distance of nuclear pixels to the nearest speckle in the same nucleus (mean ± SEM across loci, S to 180 min). In dot plots, points are individual TAS and bars are medians, except in f,l, where points are active nuclei and bars are means. In c, high and low groups were compared by linear mixed models with Holm adjustment. Two-sided Wilcoxon rank-sum tests compared each GapmeR with time-matched CTRL (d-f,l-n,q) or high with low preFOSIL quarters (h,i,k; *FOS* exon signal in i). In g,j, two-sided Fisher’s exact tests compared proximal fractions and two-sample Kolmogorov-Smirnov tests compared all distances with CTRL. P values outside c are unadjusted (Methods). *P < 0.05; **P < 0.01; ***P < 0.001; ****P < 0.0001; ns, not significant. See also Figure S5.

In WT untransfected cells, we noticed that while preFOS and *FOS* signal is usually consistently observed at the five sites, only a subset of those showed co-expression of *FOSIL*, suggesting a baseline heterogeneity of *FOSIL* transcription. Indeed, the intensity of the preFOSIL HCR FISH signal readily divided the TASes into an ‘active’ and ‘silent’ group. In unperturbed cells higher preFOSIL signal correlated with greater *FOS* exon and preFOS signal at 15 and 30 min (Fig. 5c). This correlation was observed both across nuclei and within individual nuclei (Fig. S5b).

Treatment with the L1 and L2 GapmeRs lowered preFOSIL at every time point, as expected (Fig. 5d,n). Consistent with our qPCR data, across the three conditions and three time points, the *FOS* exon signal was lower with L1 at 15 min but higher with both GapmeRs at 30 and 60 min, reflecting a prolonged and delayed induction of FOS in cells where FOSIL was perturbed (Fig. 5e,m). At 60 min, the fraction of nuclei with detectable FOS transcription was significantly higher, and often had multiple *FOS*-expressing loci (25.6% of active nuclei in control vs. 49.7% and 44.0% for L1 and L2, respectively; P = 0.0030 and P = 0.024) (Fig. 5f), a result consistent across separately imaged cells (Fig. 5l).

We then performed a second set of experiments (M-series) in which we measured *FOS* exon and preFOSIL signals, together with probes for the MALAT1 lncRNA, which is enriched at the speckle periphery ^25^ (Fig. 5g,j). Here, FOS TASes were detected using just the *FOS* exon foci which required manual TAS calling of 25% of the images to better define the segmentation thresholds. To score proximity, we used an operational cutoff of 0.32 µm, comparable to the mean distances reported for TSA-seq-defined speckle-associated domains, genomic regions consistently positioned near speckles ^58,59^. In control GapmeR-treated cells, median *MALAT1* area ranged from 1.0-to 1.3-fold relative to serum across the time course, suggesting no major speckle reorganization (Fig. S5a). At 15 min, the fraction of detected active sites within 2D-projected distance of 0.32 µm of MALAT1-defined nuclear speckles was 56.7% in CTRL against 47.7% in L1 and 38.4% in L2 (P = 2 × 10⁻⁴ and 9.31 × 10⁻¹⁴, respectively). The same direction was evident within 0.15 µm. Proximal fractions remained lower after depletion at 30 min, 41.2% in CTRL against 27.2% in L1 and 31.7% in L2. In the M series, the distributions of distances for *FOS* exon-detected sites differed from CTRL for both GapmeRs at 15 and 30 min and for L2 at 60 min (Fig. S5c-e). These fractions use independently detected active-site populations from separate imaging series (Fig. 5a). The active sites with the most preFOSIL lay nearest to speckles at 15 and 30 min (Fig. 5h,k), and at 15 min carried more *FOS* exon signal at every distance (Fig. 5i). In CTRL at 15 min, the median distance was 0.219 µm for the top preFOSIL quarter against 0.380 µm for the bottom. These high-versus-low relationships persisted within the L1 and L2 conditions after depletion.

To elucidate whether the association to speckles preceded induction or was maintained at later timepoints when the absence of transcription does not allow us to use TAS as readout for loci location, we used DNA immunoFISH. We combined a *FOS*-region DNA probe that includes the *FOSIL* locus and the upstream region of *FOS* (Fig. 5o) with SC35 immunostaining and measured normalized locus-to-speckle distances through HRG induction by DNA immunoFISH (Fig. 5o-q). This allowed us to examine locus position at all time points, including S, 0 and 180 min, albeit without a readout of RNA content. In CTRL-treated cells, distances increased slightly upon starvation, decreased sharply at 15 and 30 min after HRG induction, and subsequently returned to baseline (Fig. 5q). In GapmeR-treated cells, the locus was positioned slightly closer to speckles than in CTRL at starvation (0 min) and showed a blunted approach to speckles after HRG, remaining significantly further than CTRL at 15 and 30 min (Fig. 5q), matching the post-induction results observed by RNA smFISH. A similar trend was observed in the percentage of speckle-distant (>0.32 µm) loci (Fig. S5l), whereas the opposite was found at loci overlapping speckles (Fig. S5i) or inside the speckle-associated compartments (Fig. S5j,k), where *FOSIL* depletion led to a reduced speckle-proximal fraction mainly at 15 and 30 minutes after HRG treatment.

In summary, we show that *FOSIL*-depletion affects local folding in starved cells prior to induction (Fig. 4) and speckle positioning largely after induction (Fig. 5). Together, these results suggest that *FOSIL* transcription helps establish a baseline conformational arrangement that sustains fast and timely response to HRG, including association to nuclear speckles of the active *FOS* locus and the timely decline of its response.

### Endogenous *FOSIL* activation increases *FOS* output and speckle-proximal association of active loci

To test whether increasing endogenous *FOSIL* production could enhance *FOS* expression, we targeted its promoter with CRISPR activation (CRISPRa). We used the SAM system in MCF7 cells stably expressing dCas9-VP64 and MS2-p65-HSF1, with transient transfection of MS2-containing guide RNAs ^60^. We used a mixture of two guides to target the *FOSIL* promoter. Cells were transfected 24 hours prior to serum starvation and HRG stimulation. *FOSIL* RNA was higher in cells carrying the guides (FOSILa) than in empty guide controls throughout the RT-qPCR time course (1.7-to 4.0-fold; Fig. 6a). *FOS* mRNA was lower in FOSILa cells in serum and after starvation, but reached 3.8, 3.0 and 2.3 times the control levels at 15, 30 and 60 min after HRG, respectively. Similar trends were observed by RNA-seq (Fig. 6a,b).

**Figure 6.**
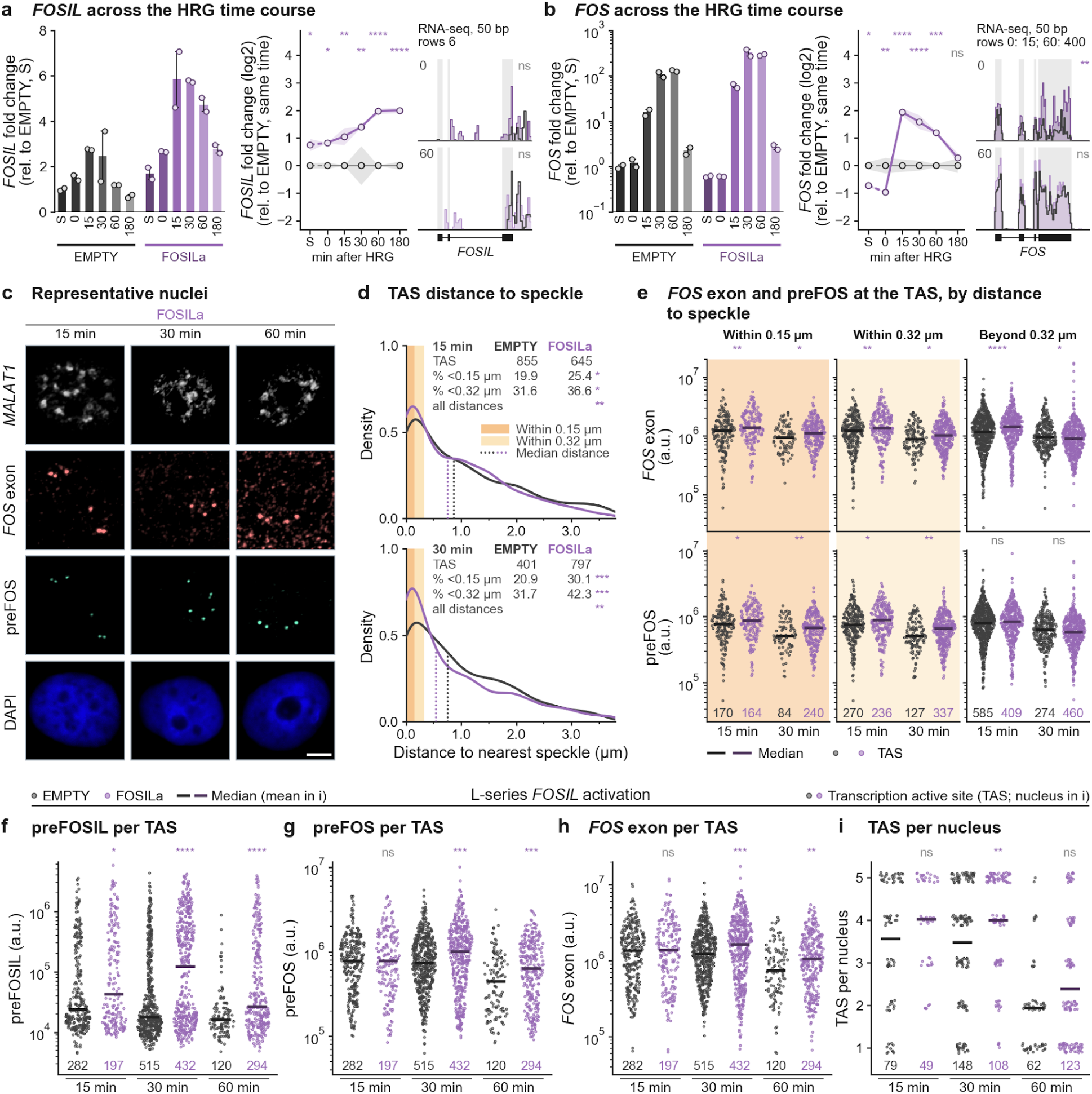
Endogenous *FOSIL* activation increases *FOS* output and speckle-proximal association of active loci. **(a,b)** *FOSIL* (a) and *FOS* (b) expression across the HRG time course in cells transfected with CRISPRa guides targeting the *FOSIL* promoter (FOSILa) or an empty guide control (EMPTY). Left, *ACTB*-normalized RT-qPCR relative to EMPTY in serum (S; mean ± SEM, two biological replicates). Middle, log2 fold change relative to time-matched EMPTY. Right, mean poly(A)+ RNA-seq coverage in 50-bp bins at 0 and 60 min (three libraries per condition and time, except two for EMPTY at 60 min). **(c)** Representative FOSILa nuclei imaged for *MALAT1*, *FOS* exon, preFOS and DAPI at 15, 30 and 60 min after HRG. Scale bar, 5 µm. The *MALAT1* display threshold was raised at 30 and 60 min for the images only. **(d)** Two-dimensional distance from each transcription active site (TAS) to the nearest *MALAT1*-defined speckle at 15 and 30 min in EMPTY and FOSILa cells. Dotted lines mark medians; tables give site counts and percentages within 0.15 and 0.32 µm (shaded). **(e)** Integrated *FOS* exon and preFOS intensity per TAS by speckle distance at 15 and 30 min (within 0.15 µm, within 0.32 µm cumulative, or beyond 0.32 µm). Points show TAS and bars show medians. **(f-i)** L-series RNA-HCR in EMPTY and FOSILa cells at 15, 30 and 60 min, in nuclei with one to five TAS. Panels f-h show preFOSIL, preFOS and *FOS* exon intensity per TAS, respectively (bars, medians); i shows TAS per active nucleus (bars, means). In a,b, qPCR comparisons used pooled-variance t-tests on ΔCt; RNA-seq comparisons used DESeq2 Wald tests with Benjamini-Hochberg adjustment across the two genes and two time points. FOSILa was compared with time-matched EMPTY by two-sided Fisher’s exact tests for proximal fractions (d) and two-sided Wilcoxon rank-sum tests for all distances (d), per-site intensities (e-h) and TAS per active nucleus (i). *P < 0.05; **P < 0.01; ***P < 0.001; ****P < 0.0001; ns, not significant.

After *FOSIL* activation, the fraction of detected active *FOS* loci within 0.32 µm of a *MALAT1*-marked speckle increased from 31.6% in the empty guide control to 36.6% at 15 min, and the fraction within 0.15 µm from 19.9% to 25.4% (Fig. 6c,d). The difference persisted and increased at 30 min. These within-series comparisons show enrichment of speckle-proximal active loci at both early time points. *FOS* exon and preFOS signals were higher after activation at both speckle-proximal compartments (Fig. 6e). Beyond this distance, *FOS* exon signal was higher at 15 min but slightly lower at 30 min, whereas preFOS signal did not differ significantly.

In separate L-series RNA-HCR experiments, FOSILa increased preFOSIL signal per active site at 15, 30 and 60 min (Fig. 6f). Both preFOS and *FOS* exon signals were higher at 30 and 60 min, and more loci per nucleus were active at 30 min (Fig. 6g-i).

Endogenous *FOSIL* activation thus increased *FOS* output and increased the speckle-proximal fraction of active loci. Its effect contrasts with ectopic *FOSIL* expression, which did not enhance induction in wild-type cells (Fig. 2k). This contrast supports a regulatory contribution of the endogenous locus and its transcriptional and processing state.

## Discussion

*FOS* mRNA is induced transiently by many stimuli, yet the downstream consequences differ between them ^4,5^, so the regulation that matters is often in the shape of the induction wave. Our findings identify *FOSIL* as a regulator of the timing, magnitude and resolution of *FOS* induction, acting through the endogenous locus rather than through an autonomous mature RNA product. Our results place local organization and nascent RNA transcription or co-transcriptional processing at the center of the mechanism. This model fits the evolutionary pattern of *FOSIL*, where strong sequence constraint at the promoter SRE and exon-3 CTCF motifs and conserved mammalian transcription coincide with rapid divergence of the exonic sequence (Fig. 1c). Moreover, the *FOSIL* SRE carries stronger SRF and ELK1 occupancy in MCF7 cells than the *FOS* promoter SRE itself (Fig. 1c), indicating that *FOSIL* receives the same immediate-early signaling input that drives *FOS*.

Studies on other *cis*-acting lncRNAs have linked chromatin conformation or subnuclear repositioning to the inducibility of neighboring genes across diverse biological timescales. For example, deletion of the *Silc1* lncRNA, located within a large TAD near *Sox11*, impaired *Sox11* induction upon peripheral nerve injury and during hippocampal responses to a novel environment, delaying axonal regeneration and memory formation ^61,62^; like *FOSIL*, *Silc1* is conserved in syntenic position and promoter elements despite divergent exonic sequence. Similarly, transcription of *ThymoD* promotes timely *Bcl11b* activation during T cell commitment by releasing the locus from the nuclear lamina and facilitating enhancer-promoter folding ^63^. In these cases, where *cis*-acting lncRNAs have been studied *in vivo*, delays in target gene induction manifest on the scale of hours to days, making it difficult to resolve the immediate molecular consequences of noncoding transcription. In cultured human cells, nascent transcription of the lncRNA *AMANZI* is important to form a chromatin loop that induces *IL37* and represses *IL1B* over 12 to 24 h of LPS stimulation in monocytes ^15^, while at shorter timescales *PACER* promotes *PTGS2* (*COX2*) induction within 2 h of LPS treatment, without a known effect on chromatin conformation, which is difficult to study for *PACER* because it is divergently transcribed from *COX2* ^64^. Across these systems, perturbing a noncoding locus effectively altered target gene induction dynamics; however, when the lncRNA acts relative to stimulus arrival and how its activity is coupled to rapid changes in chromatin folding and subnuclear positioning have remained unclear.

By tracking the *FOS*-*FOSIL* locus at minute resolution, we find that *FOSIL* perturbations act in a temporal sequence that precedes the main *FOS* mRNA and protein wave. In control cells, total *FOSIL* RNA rises about fivefold during serum starvation and peaks 15 min after HRG stimulation, ahead of the 30-min peak of *FOS* mRNA (Fig. 2c,g). Correspondingly, depleting *FOSIL* alters local chromatin compaction already in starved cells before HRG is added (Fig. 4) and blunts nuclear speckle association within 15 to 30 min of stimulation (Fig. 5). This is in contrast to the effects on bulk *FOS* mRNA persistence and FOS protein accumulation, which manifest between 60 and 180 min (Fig. 2g-j). Because enhancer and promoter-proximal noncoding RNA transcription broadly precedes activation of adjacent protein-coding genes across stimulus-responsive programs ^65^, we suggest that early noncoding transcription may serve a general preparatory role by establishing local chromatin arrangements tuned for timely target induction.

Our Capture Micro-C maps separate this normal preparatory remodeling from its disruption upon *FOSIL* loss. In unperturbed cells, whether fed or serum-starved, the *FOS*-*FOSIL* neighborhood is pre-assembled within a CTCF-flanked domain in which *FOSIL* is the most prominent contact hub (Fig. 3a,b; Fig. S3d). Serum starvation and HRG stimulation do not create or disassemble loop domains, but instead selectively redistribute contacts among existing anchors (Fig. 3e). In particular, transitioning from serum to starvation in control cells strengthens the internal *FOS*|*FOSIL* boundary as basal *FOSIL* expression rises, increasing insulation between the two genes prior to HRG treatment (Fig. 3f,g). When *FOSIL* is depleted, architectural changes are strictly confined to the regulatory domain between upstream enhancer e1 and the CTCF_R_ anchor just downstream of *FOSIL* (Fig. 4). Specifically, *FOSIL* depletion prevents the starvation-induced strengthening of the *FOS*|*FOSIL* boundary, leaving it significantly weaker than in control cells at 0 and 15 min, while increasing local contact density across both the →*FOS* gene and the *FOS*→*FOSIL* intergenic interval from starvation onward (Fig. 4c-e). At the same time, excess contacts of the *FOS* promoter with e2, the *FOSIL*_TSS_ and the *FOSIL* exon-3 CTCF site reach their maximum in starved cells before HRG stimulation (Fig. 4a,b). Active transcription across *FOSIL* provides a plausible mechanism for both effects, as elongating RNA polymerase can generate local domain insulation ^18^, whereas reducing elongation through an intragenic CTCF site can cause cohesin to accumulate at that anchor and compact the surrounding chromatin ^66^. Notably, because this more compacted, less insulated state at 0 min is followed by a delayed *FOS* induction wave, elevated contact frequency alone is insufficient for rapid activation; instead, *FOSIL* activity appears to maintain a selective, locally insulated conformation poised for synchronous firing.

Upon HRG induction, this pre-induction chromatin state gives way to a dynamic spatial phenotype at nuclear speckles. In control cells, the *FOS* locus moves rapidly toward speckles at 15 and 30 min and returns toward baseline by 60 min (Fig. 5o-q; Fig. S5i-l), whereas *FOSIL* depletion blunts this early repositioning near speckles as seen both by DNA-FISH and smRNA-FISH assays (Fig. 5g,j,q; Fig. S5c-e,i-l). Across unperturbed and depleted cells alike, transcription active sites with higher nascent *FOSIL* signal are positioned closer to speckles and produce more *FOS* RNA (Fig. 5c,h,i,k; Fig. S5b). Conversely, endogenous CRISPR activation of the *FOSIL* promoter increases both *FOS* transcriptional output and the speckle-proximal fraction of active loci after HRG (Fig. 6a,b,d-i). However, CRISPRa transactivators have been recently described to promote speckle association and may contribute to this gain-of-function phenotype ^30^.

While local chromatin compaction and speckle association operate at different physical and time scales, recent work indicates that they can be functionally linked; for example, depleting speckle scaffolding proteins alters contacts within active compartments ^67^, in turn, loss of CTCF or cohesin compromises gene-speckle association ^31^, although these are likely context dependent as it has also been described that most chromatin loops survive acute speckle disruption ^68^. At the active locus, speckle proximity has been shown to amplify transcriptional bursts at stress- and p53-responsive genes^24,69^, enhance co-transcriptional splicing efficiency^23^, and modulate *FOS* pre-mRNA splicing during ribotoxic stress^70^. At *FOSIL*, these observations raise the possibility that co-transcriptional assembly of the nascent *FOSIL* ribonucleoprotein complex contributes to speckle engagement before a mature transcript is released. Perturbing splice-site recognition with RNase-H-inactive, splice-occluding oligonucleotides at the endogenous locus will provide a direct test of this processing contribution without degrading the *FOSIL* transcript.

Beyond the initial burst, the delayed bulk *FOS* mRNA and protein wave (Fig. 2g-j) and the persistence of active transcription sites at 60 min (Fig. 5e,f,l,m; Fig. S5f-h) reveal that *FOSIL* regulation extends to also shape the resolution of the immediate-early response. In *FOSIL*-depleted cells, reduced *FOS* output near speckles at 15 to 30 min is followed at 60 min by an increased fraction of nuclei retaining multiple active *FOS* alleles and elevated *FOS* exon signal at speckle-distant sites (Fig. 5f,l; Fig. S5f-h). This shift from a sharp, speckle-proximal burst at 15 to 30 min to prolonged, speckle-distant firing at 60 min suggests that without *FOSIL*, individual alleles activate less synchronously and remain transcriptionally active for longer. Together with the absence of an effect from ectopic *FOSIL* expression (Fig. 2k), these findings point to the *cis* transcriptional and co-transcriptional state of the endogenous locus as the driver of coordinated induction and timely shutoff.

Immediate-early genes must translate extracellular cues into precisely shaped transcriptional pulses whose amplitude and duration guide downstream cell-fate decisions ^4,5^. Whereas classical models of *FOS* induction have focused on signal-dependent transcription factor modification at the promoter and upstream enhancers, our study identifies an adjacent, co-induced lncRNA locus as a kinetic organizer of the response. In our working model, basal and early *FOSIL* transcription restrains local chromatin compaction and maintains domain insulation prior to the arrival of a stimulus, enabling rapid, synchronous engagement of the activated *FOS* locus with the nuclear speckle periphery and timely resolution of the transcriptional pulse. Because thousands of mammalian stimulus-responsive genes are paired with inducible noncoding transcription units that fire ahead of the coding burst ^10,11,65^, coupling preparatory 3D chromatin organization to dynamic subnuclear positioning may represent a general principle by which noncoding loci fine-tune the speed and precision of inducible gene expression.

### Limitations of the study

RNase-H-active GapmeRs can affect elongation as well as RNA abundance ^46^. GapmeR cleavage of the nascent transcript also removes splicing and 3′ end formation along with the RNA, so the transcript, its processing and the act of transcribing it are not separated here. Finally, ectopic expression supplies the mature transcript *in trans* and does not recapitulate endogenous co-transcriptional production or local chromatin context.

## Materials and methods

## Resource availability

### Lead contact

Further information and requests for resources and reagents should be directed to Igor Ulitsky.

### Materials availability

Materials generated in this study are available from the lead contact upon reasonable request.

### Data and code availability

Analysis code and additional data supporting the findings are available from the lead contact upon reasonable request.

## Experimental model details

### Cell lines

Experiments were performed in human MCF7 breast-cancer cells (single-cell-derived WT line and a derivative stably expressing dCas9-VP64 and MS2-p65-HSF1 for CRISPRa). Cells were maintained in DMEM containing 4.5 g/L glucose, 4 mM L-glutamine, 10% fetal bovine serum (FBS) and 1% penicillin-streptomycin at 37°C and 5% CO_2_.

## Method details

### Serum starvation and stimulation

For all transfection experiments, medium was replaced with serum-free basal medium 24 h after transfection, and cells were serum-starved for 16-20 h before stimulation in complete medium containing 10% FBS and 100 ng/mL recombinant human NRG1-β1/HRG1-β1 EGF domain (R&D Systems, 396-HB-050/CF).

### *FOSIL* depletion with LNA-GapmeRs

LNA-GapmeRs (Qiagen) were used at 50 nM final concentration with sequences (5′-3′) AACACGTCTATACGC (CTRL), TCCTCAGAAAGAAACGAAAG (L1) and ACAGCGTGTAAGAGGAAGTTGG (L2). For expression assays, 200,000 cells were seeded per well of a 12-well plate the day before forward transfection with 2 µL Lipofectamine 3000 per well. Imaging experiments used reverse transfection in chambered slides. For large-format reverse transfection used in Capture Micro-C, 1 nmol GapmeR and 75 µL Lipofectamine 3000 were diluted separately in 1.5 mL Opti-MEM, incubated together for 15 min at room temperature, and added to a 15-cm dish before seeding 15 million MCF7 cells in 17 mL medium containing 10% FBS.

### Endogenous CRISPR activation

MCF7 cells stably expressing dCas9-VP64 and MS2-p65-HSF1 were transiently transfected with MS2-containing sgRNA plasmids using linear 25-kDa PEI (Polysciences, 23966) at 1 µg total DNA per 12-well well (and proportionally scaled for imaging), complexed in 150 mM NaCl for 20 min at room temperature. *FOSIL* activation (FOSILa) used an equimolar mix of two guides (TCCCGCCCACAGCCTACCC and GCCCACAGCCTACCCCGGG; 500 ng each per 12-well well). Control cells received the empty guide vector (EMPTY).

### Ectopic *FOSIL* expression

The four-exon *FOSIL* cDNA sequence (*LINC01220*-203, synthesized by Twist Bioscience) was cloned into pcDNA3.1(+) under the CMV promoter. WT cells were transfected with 500 ng *FOSIL* plasmid plus 500 ng empty pcDNA3.1(+), or 1 µg empty vector, using PEI and the starvation and stimulation schedule above.

### Proliferation assay

Cell proliferation was measured with PrestoBlue on a GloMax Explorer (Promega; excitation 520 nm, emission 580-640 nm). Cells were cotransfected using Lipofectamine 3000 with empty or *FOSIL* plasmid together with pooled L1 + L2 GapmeRs (50 nM each) or CTRL GapmeR matched at 100 nM total concentration, followed by serum starvation and HRG stimulation. Separate wells were read at 12 and 24 h (two wells per condition in each of two independent experiments).

### RNA quantification and reverse transcription qPCR

Total RNA was isolated with TRIzol, quantified by NanoDrop spectrophotometry, and treated with Baseline-ZERO DNase for 10 min at 37°C. One microgram of RNA was reverse-transcribed using the qScript Flex cDNA Synthesis Kit (Quantabio) with mixed oligo(dT) and random primers. qPCR was performed with Fast SYBR Green Master Mix (Applied Biosystems) on a ViiA 7 instrument in 384-well plates in technical triplicate. Relative abundance was calculated as 2^−ΔCt^ normalized to *ACTB* or *GAPDH*.

### Flow cytometry of FOS protein

Trypsinized cells were washed twice in FACS buffer (PBS with 4% FBS and 2 mM EDTA), and up to 1 million cells in 100 µL were fixed with 0.5 mL cold 4% PFA in PBS for 10 min at room temperature (three separately transfected cultures pooled per condition). Cells were washed twice, permeabilized and stained for 30 min at room temperature in the dark in 200 µL PBS containing 0.1% Triton X-100, 4% FBS, 2 mM EDTA and 1 µL PE-conjugated c-Fos (9F6) rabbit monoclonal antibody (Cell Signaling Technology, 14609), then washed and resuspended in FACS buffer. Single cells and FOS-positive gates were set in FlowJo using stained and unstained WT controls stimulated with HRG for 1 h.

### Multiplex RNA HCR

Multiplex RNA smFISH was performed using HCR v3.0 ^71^ on chambered slides. Cells were fixed in 4% formaldehyde for 10 min at room temperature, washed in DPBS, and permeabilized in ice-cold 70% ethanol overnight at −20°C. Samples were rinsed twice in 2× SSC and once in 5× SSCT, prehybridized for 30 min at 37°C, and hybridized overnight at 37°C in HCR hybridization buffer supplemented with salmon-sperm DNA using probe sets targeting *FOSIL* introns (HSS-LINC01220introns-B4), *FOS* introns (B2), *FOS* exons (B5) and *MALAT1* (B2). Slides were washed four to five times for 5 min at 37°C in HCR wash buffer (30% formamide, 5× SSC, 9 mM citric acid, 0.1% Tween-20, 50 µg/mL heparin) and twice for 5 min in 5× SSCT at room temperature. After 30 min preamplification, snap-cooled hairpins (B4-Alexa Fluor 647, B2-Alexa Fluor 488, B5-Alexa Fluor 546; 60 nM each) were incubated for 4 h at room temperature in the dark, followed by five 5-min washes in 5× SSCT, DAPI counterstaining (1:2000, 5 min) and mounting.

### DNA immuno-FISH for *FOS* locus and speckle positioning

DNA immuno-FISH was adapted from Potapova et al. ^72^. Cells on chamber slides or #1.5 coverslips were fixed in 4% PFA for 15 min, permeabilized in 0.1% Triton X-100/DPBS for 15 min, and treated with RNase A (2 µg/mL, 30 min, 37°C). Samples underwent two liquid-nitrogen freeze-thaw cycles, 0.1 N HCl treatment for 5 min, 2× SSC washes, and 30 min preincubation in 50% formamide/2× SSC at 37°C. The *FOS* locus probe (Empire Genomics, FOS-20-OR; 0.4 µL in 4.6 µL hybridization buffer) was sealed under a HybriSlip with Cytobond, codenatured with the sample at 85°C for 7 min, and hybridized for 48-72 h at 37°C.

Post-hybridization washes were three 10-min washes in 50% formamide/2× SSC at 45°C, two 5-min washes in 1× SSC at 45°C, and one 5-min wash in 1× SSC at room temperature. Samples were blocked for 1 h in 5% BSA/0.5% Triton X-100/DPBS, incubated with mouse anti-SC35 (Abcam, ab11826; 1:100, 2 h) and goat anti-mouse Alexa Fluor 488 (Abcam, ab150113; 1:500, 1 h) in 2.5% BSA/0.25% Triton X-100/DPBS, counterstained with DAPI (1:1000, 5 min), and mounted in ProLong Diamond.

### Microscopy acquisition

Images were acquired on a Nikon Ti2 spinning-disk confocal microscope (Yokogawa CSU-W1, Prime BSI camera, Plan Apo λ 60×/1.4 NA oil objective) at 0.1083 µm per pixel with 0.2-µm z-steps and analyzed as 2D projections.

### Capture Micro-C library preparation and target enrichment

Capture Micro-C was adapted from the Micro Capture-C workflow ^21,73,74^. Aliquots of 5 million MCF7 cells were fixed in 2% PFA for 10 min, quenched with 0.15 M glycine, permeabilized in 0.006% digitonin for 10 min, immobilized on concanavalin A beads (EpiCypher, 21-1411), and digested with 500 gel units MNase (NEB, M0247) for 1 h at 37°C. End repair, biotin labeling, proximity ligation, crosslink reversal and DNA purification followed the standard Micro Capture-C protocol ^73,74^. Purified DNA was sonicated (Diagenode Bioruptor) to 180-200 bp, enriched on streptavidin beads, ligated to adapters (NEBNext Ultra II), and indexed on-bead using KAPA HiFi HotStart ReadyMix (7-8 cycles). Target enrichment used custom Twist 70-mer capture probes spanning the *FOS* neighborhood (design TE-95053285; Twist Standard Hybridization v2 protocol). Libraries were pooled 12 per capture reaction at 166.7 ng each (2,000 ng total), with each of the four GapmeR capture reactions containing one complete biological replicate of CTRL, L1 and L2 across all four time points (12 post-capture PCR cycles), and sequenced paired-end (2 × 150 bp) on an Illumina NovaSeq X Plus.

### RNA-seq library preparation

Total RNA-seq libraries for the GapmeR time course (two independent biological replicates per condition at S, 0, 15, 30 and 60 min) were prepared using the NEBNext rRNA Depletion Kit v2 and NEBNext Ultra II Directional RNA Library Prep Kit; poly(A)+ RNA-seq libraries for CRISPRa were prepared using the Lexogen Poly(A) RNA Selection and CORALL mRNA-Seq Library Prep Kits. Libraries were sequenced paired-end (2 × 61 bp).

### Computational and statistical analyses

Key resources, computational processing of Capture Micro-C and RNA-seq data, public dataset integration, evolutionary sequence conservation and motif analyses, image segmentation, and statistical testing are detailed in the Supplementary Methods.

## Supporting information

FOSIL_Supplementary_Methods

---

**Figure S1.**
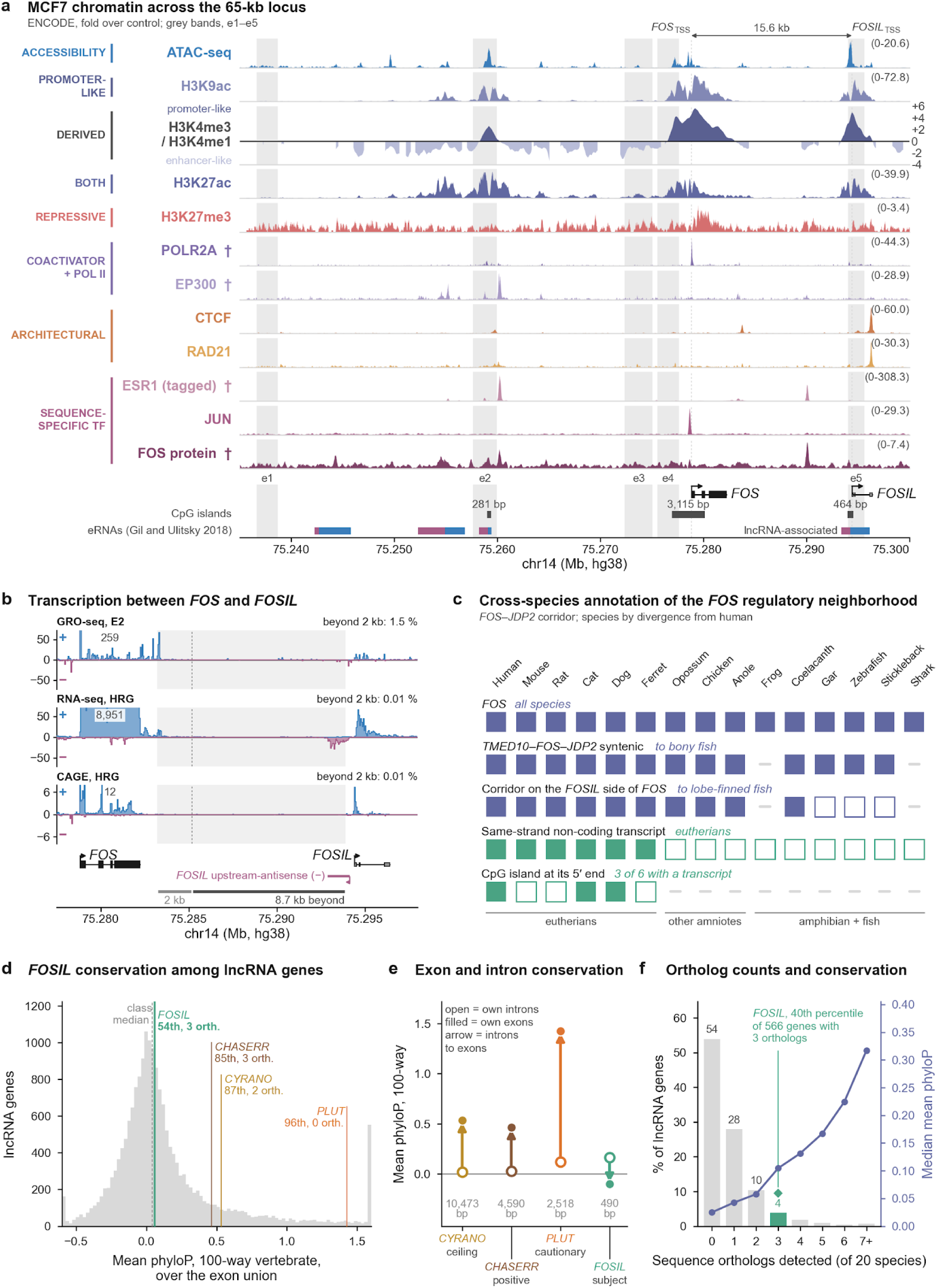
The *FOSIL* locus in MCF7 cells and across species, related to Figure 1. **(a)** MCF7 chromatin and transcription factor profiles across the 65-kb *FOS*-*FOSIL* region (GRCh38), with candidate regulatory intervals e1-e5 (grey bands), CpG islands and MCF7 GRO-seq enhancer RNAs (eRNAs, plus strand in blue and minus strand in plum) ^44,45^. Daggers mark unspecified POLR2A/EP300 growth conditions, tagged ESR1 and FOS protein ChIP. **(b)** Plus-strand (above) and minus-strand (below) estradiol GRO-seq, HRG total RNA-seq (CTRL) and HRG CAGE pooled across time points in 50-bp bins (chr14:75,277,500-75,298,000). Percentages give intergenic plus-strand signal per kb, starting 2 kb downstream of *FOS*, relative to the *FOS* gene body. Minus-strand signal reflects upstream antisense transcription at *FOSIL*. **(c)** Gene order, same-strand noncoding transcripts and their 5′ CpG islands in the *TMED10*-*FOS*-*JDP2* region across 15 vertebrates (filled and open squares, present or absent in the annotation, respectively). Dashes indicate cases that could not be assessed. **(d)** Mean 100-way vertebrate phyloP across exon unions of 18,534 GENCODE v42 lncRNAs (dashed line, class median), with percentiles and sequence ortholog counts for *FOSIL* and three benchmark lncRNAs. **(e)** Mean vertebrate phyloP for each gene’s exon and intron unions (NR_038421.1 for *FOSIL*). **(f)** Percentage of 14,556 GENCODE lncRNAs in each sequence ortholog count bin (bars) and median phyloP per bin (line). The diamond marks *FOSIL*.

**Figure S2.**
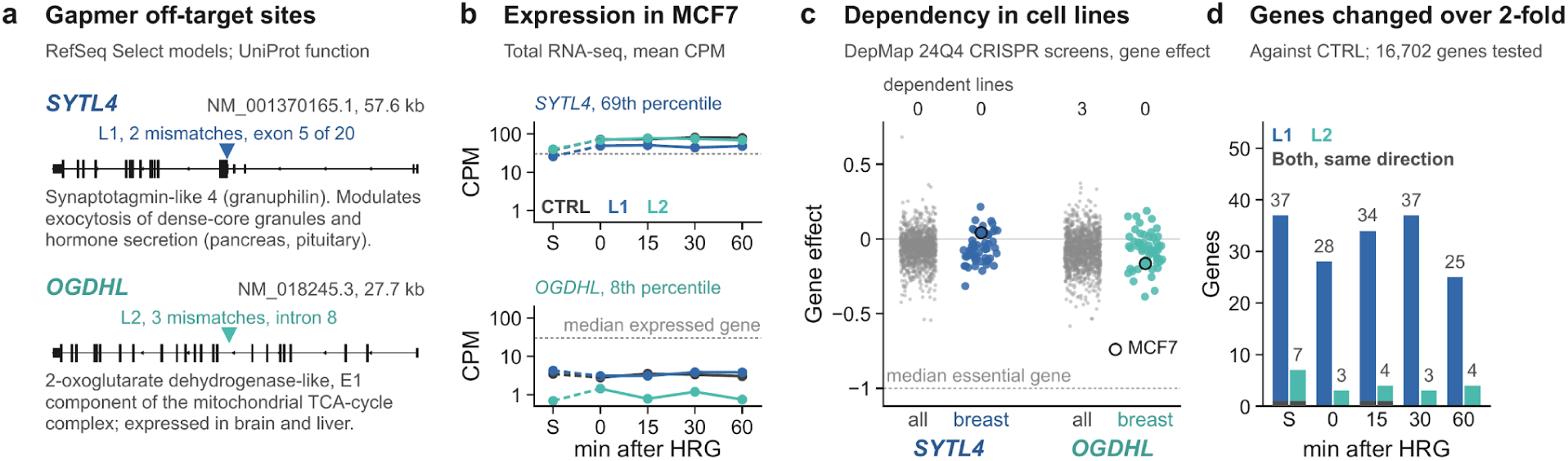
Specificity of the two *FOSIL* GapmeRs, related to Figure 2. **(a)** Candidate sense-strand off-target sites (triangles) for L1 in *SYTL4* (two mismatches) and L2 in *OGDHL* (three mismatches). **(b)** *SYTL4* and *OGDHL* expression in CTRL, L1 and L2 cells (mean counts per million, two biological replicates). Percentiles are relative to 12,048 expressed genes. **(c)** Chronos gene effects across all 1,178 DepMap 24Q4 cell lines and the 53 breast cancer lines (open circle, MCF7). Zero denotes no effect. Counts indicate dependent lines (dependency probability > 0.5). **(d)** Genes changing more than twofold versus time-matched CTRL (DESeq2 test for |log2FC| > 1, adjusted P < 0.05). Dark segments mark genes changed in the same direction by both GapmeRs. S denotes normal serum, 0 denotes serum starvation, and 15-60 denotes minutes after HRG.

**Figure S3.**
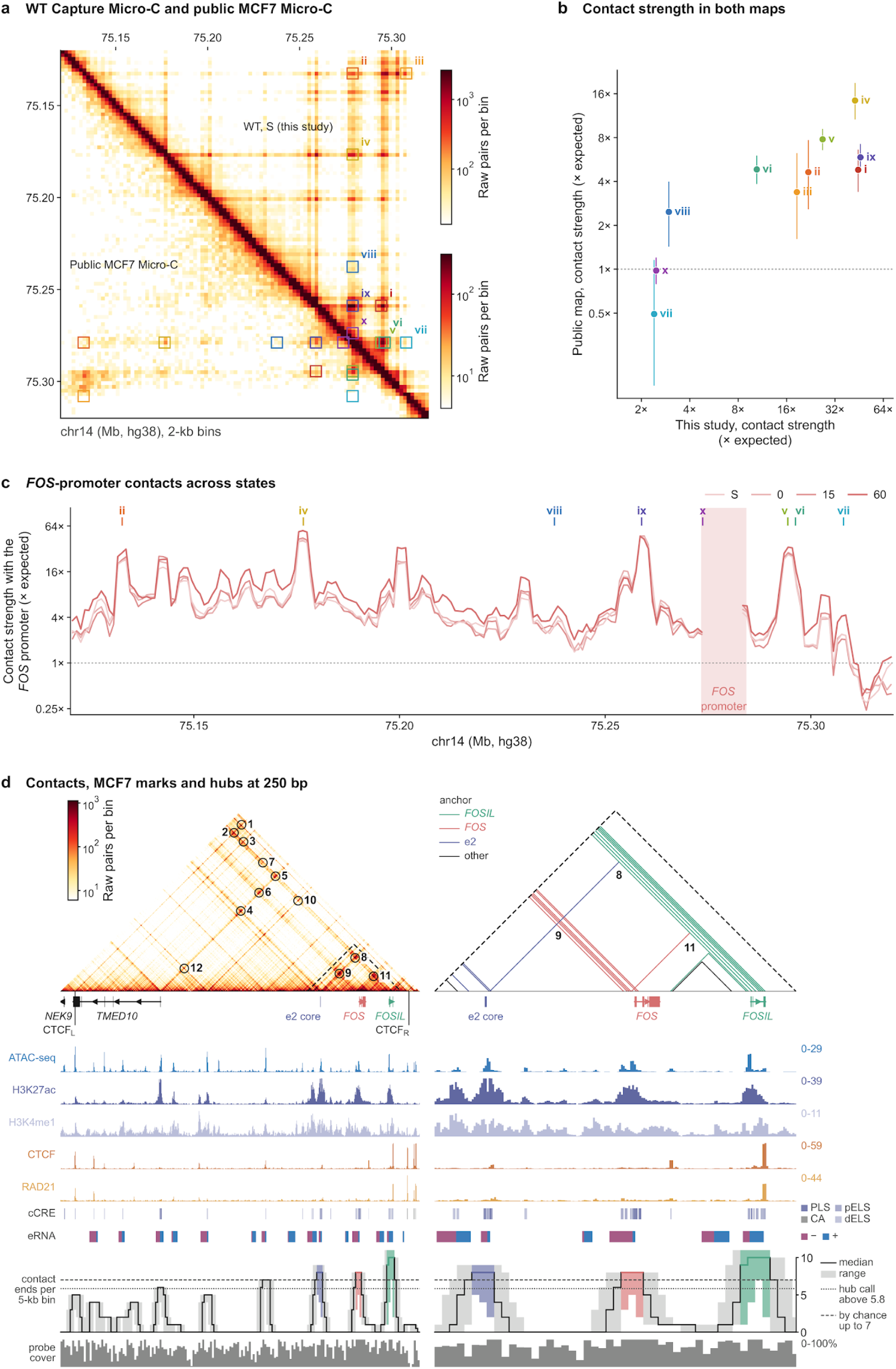
Comparison with genome-wide Micro-C and *FOS*-promoter viewpoint profiles, related to Figure 3. **(a)** The 200-kb *FOS*-*FOSIL* region at 2-kb resolution in WT Capture Micro-C (upper triangle, two serum replicates summed) and public genome-wide MCF7 Micro-C (GSE278978, lower triangle) ^52^. Each map shows unbalanced counts on its own logarithmic scale. Boxed contacts are (i) e2 × *FOSIL*_TSS_, (ii) CTCF_L_ × *FOS*, (iii) CTCF_L_ × CTCF_R_, (iv) *TMED10* × *FOS*, (v) *FOS* × *FOSIL*_TSS_, (vi) *FOS* × *FOSIL*_CTCF_, (vii) *FOS* × CTCF_R_, (viii) e1 × *FOS*, (ix) e2 × *FOS* and (x) e3 × *FOS*. **(b)** Contact strength for i-x (observed over distance-expected counts in 3 × 3-kb boxes). WT values are means of two replicates. Error bars are 95% Poisson confidence intervals for the public map. **(c)** Virtual 4C profiles with a 3 × 3-kb window anchored at the *FOS* promoter, in serum (S), after starvation (0) and at 15 and 60 min after HRG (mean of two WT replicates). Shading marks bins excluded near the viewpoint. **(d)** Pooled 250-bp map of chr14:75,125,000-75,310,000 (48 CTRL, L1 and L2 libraries). Left, contacts 1-12 from Figure 3 are circled. Right, the outlined region with contacts called by Mustache at both 250 bp and 1 kb, colored by anchor. Tracks show MCF7 ATAC-seq and ChIP-seq, ENCODE cCREs (promoter-like [PLS], proximal and distal enhancer-like [pELS, dELS], and other accessible chromatin [CA]), strand-specific GRO-seq eRNAs, contact ends per 5-kb bin and capture-probe coverage. The hub threshold is 1.5 SD above the mean, and the chance level is the 95th percentile of the maximum bin count across random placements (Methods).

**Figure S4.**
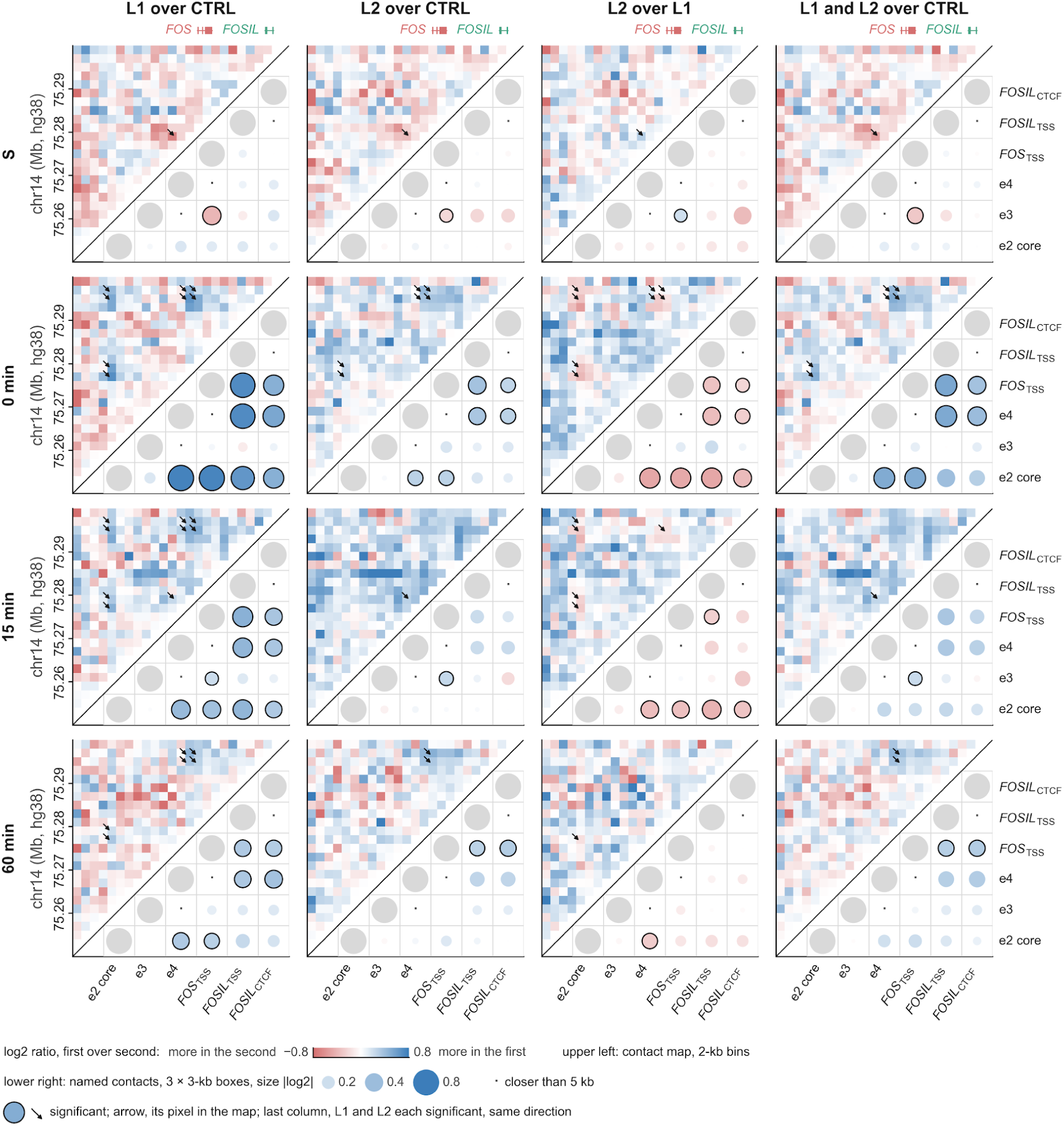
Pairwise differential contact maps and landmark interactions across states, related to Figure 4. Differential Capture Micro-C maps across chr14:75.25-75.30 Mb in normal serum (S), after starvation (0) and at 15 and 60 min after HRG (rows). Columns compare L1 with CTRL, L2 with CTRL, L2 with L1, and L1 and L2 together with CTRL. Upper-left triangles show log2 contact ratios in 2-kb bins; the fourth column shows the mean of the first two log2 ratios. Lower-right triangles show log2 fold changes in read pairs within 3 × 3-kb boxes for contacts among e2 core, e3, e4, *FOS*_TSS_, *FOSIL*_TSS_ and *FOSIL*_CTCF_. Circle area indicates magnitude; blue and red indicate higher and lower contact in the first condition, respectively. Grey discs mark the landmarks on the diagonal, and dots mark pairs less than 5 kb apart, which were not scored. Rings mark significant contacts (diffHic/edgeR, FDR < 1%) and arrows identify their pixels on the map. In the fourth column, a ring requires a significant change in the same direction for both GapmeRs (Methods).

**Figure S5.**
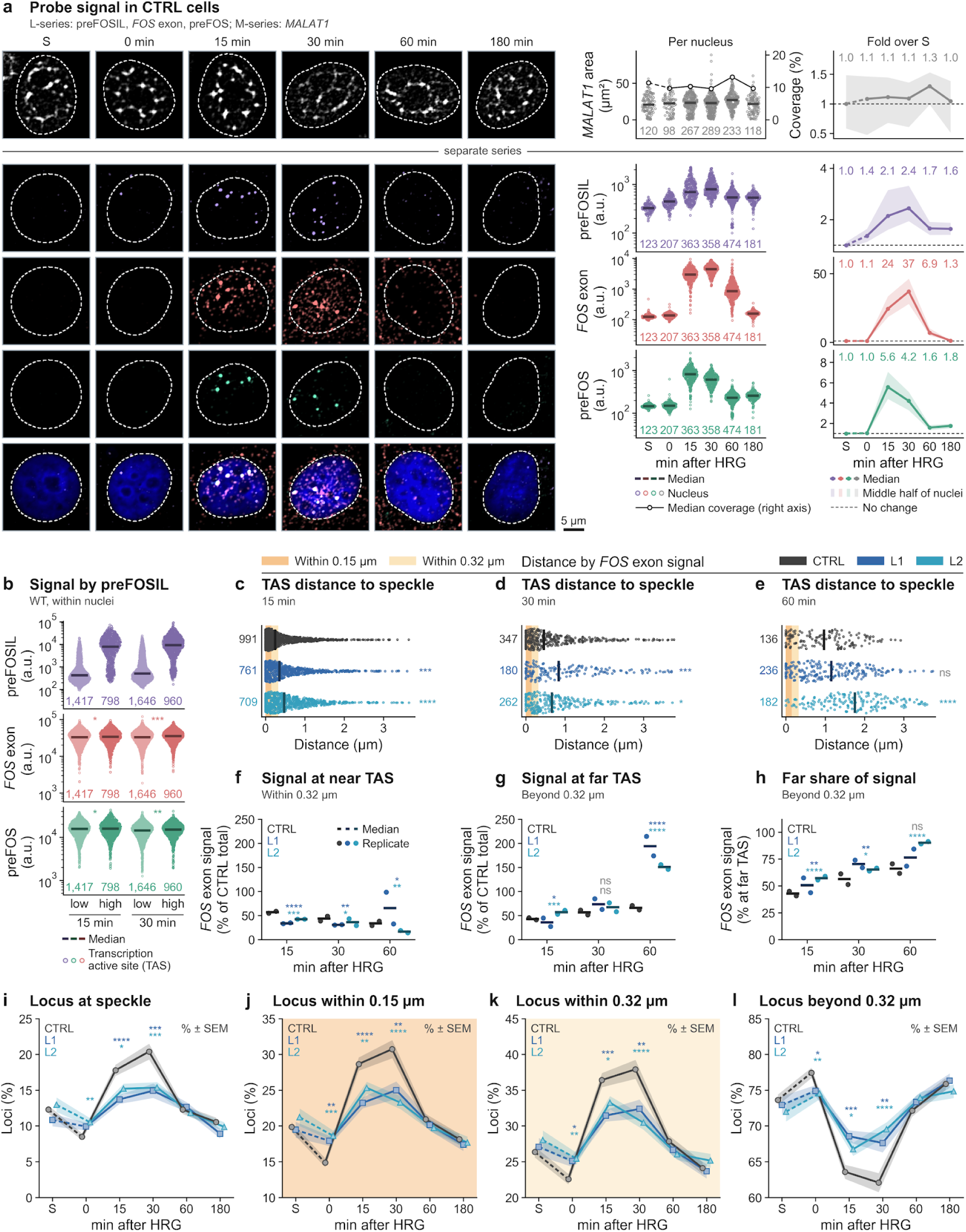
*FOS* RNA output and speckle proximity in single cells, related to Figure 5. **(a)** RNA-HCR probe specificity test in CTRL GapmeR-transfected cells from serum (S) to 180 min after HRG. The M series (*MALAT1*, top) and L series (preFOSIL, *FOS* exon and preFOS, below) were imaged separately. Left, representative nuclei near each time point’s median, shown as average-intensity projections with maximum-projected DAPI (scale bar, 5 µm). Right, *MALAT1* area and probe intensities per nucleus (bars, medians; numbers, nuclei), and fold change relative to serum. **(b)** Within-nucleus comparison of preFOS and *FOS* exon intensity per TAS in WT cells at 15 and 30 min (L series). Sites were split at the mean preFOSIL intensity of their own nucleus (at least two TAS). **(c-e)** Distance from each TAS detected by *FOS* exon signal to the nearest *MALAT1*-defined speckle at 15 (c), 30 (d) and 60 min (e) in the M series. Shading marks the distances defined in Figure 5g. **(f,g)** Integrated *FOS* exon signal summed per nucleus at TAS within (f) or beyond (g) 0.32 µm of a speckle at 15, 30 and 60 min (M series). Each point shows the replicate mean as a percentage of its time-matched CTRL total (two biological replicates; bars, medians). **(h)** Share of each condition’s total *FOS* exon signal at TAS beyond 0.32 µm at each time point. **(i-l)** Percentage of DNA-FISH loci overlapping an SC35-marked speckle (i), within 0.15 µm (j), within 0.32 µm (k) or beyond 0.32 µm (l) from S to 180 min (percentage ± SEM). The high and low groups in b were compared by linear mixed models with Holm adjustment. Comparisons with time-matched CTRL used two-sample Kolmogorov-Smirnov tests for TAS distances (c-e), two-sided Wilcoxon rank-sum tests on per-nucleus values (f-h) and two-sided Fisher’s exact tests on locus counts (i-l). Symbols and abbreviations as in Figure 5.

## References

1. Ohe, S., Kubota, Y., Yamaguchi, K., Takagi, Y., Nashimoto, J., Kozuka-Hata, H., Oyama, M., Furukawa, Y., and Takekawa, M. (2022). ERK-mediated NELF-A phosphorylation promotes transcription elongation of immediate-early genes by releasing promoter-proximal pausing of RNA polymerase II. Nat. Commun. 13, 7476.

2. Sugo, N., Atsumi, Y., and Yamamoto, N. (2025). Transcription and epigenetic factor dynamics in neuronal activity-dependent gene regulation. Trends Genet. 41, 425–436.

3. Nagashima, T., Shimodaira, H., Ide, K., Nakakuki, T., Tani, Y., Takahashi, K., Yumoto, N., and Hatakeyama, M. (2007). Quantitative transcriptional control of ErbB receptor signaling undergoes graded to biphasic response for cell differentiation. J. Biol. Chem. 282, 4045–4056.

4. Nakakuki, T., Birtwistle, M.R., Saeki, Y., Yumoto, N., Ide, K., Nagashima, T., Brusch, L., Ogunnaike, B.A., Okada-Hatakeyama, M., and Kholodenko, B.N. (2010). Ligand-specific c-Fos expression emerges from the spatiotemporal control of ErbB network dynamics. Cell 141, 884–896.

5. Mina, M., Magi, S., Jurman, G., Itoh, M., Kawaji, H., Lassmann, T., Arner, E., Forrest, A.R.R., Carninci, P., Hayashizaki, Y., et al. (2015). Promoter-level expression clustering identifies time development of transcriptional regulatory cascades initiated by ErbB receptors in breast cancer cells. Sci. Rep. 5, 11999.

6. Lavoie, H., Gagnon, J., and Therrien, M. (2020). ERK signalling: a master regulator of cell behaviour, life and fate. Nat. Rev. Mol. Cell Biol. 21, 607–632.

7. Ram, A., Murphy, D., DeCuzzi, N., Patankar, M., Hu, J., Pargett, M., and Albeck, J.G. (2023). A guide to ERK dynamics, part 2: downstream decoding. Biochem. J. 480, 1909–1928.

8. De Cesare, D., Jacquot, S., Hanauer, A., and Sassone-Corsi, P. (1998). Rsk-2 activity is necessary for epidermal growth factor-induced phosphorylation of CREB protein and transcription of c-fos gene. Proc. Natl. Acad. Sci. U. S. A. 95, 12202–12207.

9. Gschwind, A.R., Mualim, K.S., Karbalayghareh, A., Sheth, M.U., Dey, K.K., Jagoda, E., Nurtdinov, R.N., Xi, W., Tan, A.S., Galante, J., et al. (2026). An encyclopedia of human enhancer-gene regulatory interactions. Nature 657, 179–190.

10. Wang, L., Yuan, W., Gamliel, A., Ma, W., Lee, S., Tan, Y., Chen, Z., Taylor, H., Ohgi, K., Oh, S., et al. (2025). An eRNA transcription checkpoint for diverse signal-dependent enhancer activation programs. Nat. Genet. 57, 962–972.

11. Aitken, S., Magi, S., Alhendi, A.M.N., Itoh, M., Kawaji, H., Lassmann, T., Daub, C.O., Arner, E., Carninci, P., Forrest, A.R.R., et al. (2015). Transcriptional dynamics reveal critical roles for non-coding RNAs in the immediate-early response. PLoS Comput. Biol. 11, e1004217.

12. Gorbovytska, V., Kim, S.-K., Kuybu, F., Götze, M., Um, D., Kang, K., Pittroff, A., Brennecke, T., Schneider, L.-M., Leitner, A., et al. (2022). Enhancer RNAs stimulate Pol II pause release by harnessing multivalent interactions to NELF. Nat. Commun. 13, 2429.

13. Schaukowitch, K., Joo, J.-Y., Liu, X., Watts, J.K., Martinez, C., and Kim, T.-K. (2014). Enhancer RNA facilitates NELF release from immediate early genes. Mol. Cell 56, 29–42.

14. Fanucchi, S., Fok, E.T., Dalla, E., Shibayama, Y., Börner, K., Chang, E.Y., Stoychev, S., Imakaev, M., Grimm, D., Wang, K.C., et al. (2019). Immune genes are primed for robust transcription by proximal long noncoding RNAs located in nuclear compartments. Nat. Genet. 51, 138–150.

15. Fok, E.T., Moorlag, S.J.C.F.M., Negishi, Y., Groh, L.A., Dos Santos, J.C., Gräwe, C., Monge, V.V., Craenmehr, D.D.D., van Roosmalen, M., da Cunha Jolvino, D.P., et al. (2024). A chromatin-regulated biphasic circuit coordinates IL-1β-mediated inflammation. Nat. Genet. 56, 85–99.

16. Engreitz, J.M., Haines, J.E., Perez, E.M., Munson, G., Chen, J., Kane, M., McDonel, P.E., Guttman, M., and Lander, E.S. (2016). Local regulation of gene expression by lncRNA promoters, transcription and splicing. Nature 539, 452–455.

17. Beucher, A., Miguel-Escalada, I., Balboa, D., De Vas, M.G., Maestro, M.A., Garcia-Hurtado, J., Bernal, A., Gonzalez-Franco, R., Vargiu, P., Heyn, H., et al. (2022). The HASTER lncRNA promoter is a cis-acting transcriptional stabilizer of HNF1A. Nat. Cell Biol. 24, 1528–1540.

18. Koska, M., Nagano, M., Swigut, T., Boettiger, A.N., Hansen, A.S., and Wysocka, J. (2026). Promoter strength and position govern promoter competition through transcript-dependent insulation. Nat. Genet. 58, 1928–1940.

19. Unfried, J.P., and Ulitsky, I. (2022). Substoichiometric action of long noncoding RNAs. Nat. Cell Biol. 24, 608–615.

20. Karpinska, M.A., Zhu, Y., Fakhraei Ghazvini, Z., Ramasamy, S., Barbieri, M., Cao, T.B.N., Varahram, N., Aljahani, A., Lidschreiber, M., Papantonis, A., et al. (2025). CTCF depletion decouples enhancer-mediated gene activation from chromatin hub formation. Nat. Struct. Mol. Biol. 32, 1268–1281.

21. Goel, V.Y., Huseyin, M.K., and Hansen, A.S. (2023). Region Capture Micro-C reveals coalescence of enhancers and promoters into nested microcompartments. Nat. Genet. 55, 1048–1056.

22. Beagan, J.A., Pastuzyn, E.D., Fernandez, L.R., Guo, M.H., Feng, K., Titus, K.R., Chandrashekar, H., Shepherd, J.D., and Phillips-Cremins, J.E. (2020). Three-dimensional genome restructuring across timescales of activity-induced neuronal gene expression. Nat. Neurosci. 23, 707–717.

23. Bhat, P., Chow, A., Emert, B., Ettlin, O., Quinodoz, S.A., Strehle, M., Takei, Y., Burr, A., Goronzy, I.N., Chen, A.W., et al. (2024). Genome organization around nuclear speckles drives mRNA splicing efficiency. Nature 629, 1165–1173.

24. Kim, J., Venkata, N.C., Hernandez Gonzalez, G.A., Khanna, N., and Belmont, A.S. (2020). Gene expression amplification by nuclear speckle association. J. Cell Biol. 219, e201904046.

25. Fei, J., Jadaliha, M., Harmon, T.S., Li, I.T.S., Hua, B., Hao, Q., Holehouse, A.S., Reyer, M., Sun, Q., Freier, S.M., et al. (2017). Quantitative analysis of multilayer organization of proteins and RNA in nuclear speckles at super resolution. J. Cell Sci. 130, 4180–4192.

26. Song, Y.J., Shinn, M.K., Bangru, S., Wang, Y., Sun, Q., Hao, Q., Chaturvedi, P., Freier, S.M., Perez-Pinera, P., Nelson, E.R., et al. (2026). LncRNA-splicing factor condensates regulate hypoxia-responsive pre-mRNA processing near nuclear speckles. Mol. Cell 86, 1061–1080.e10.

27. Ilik, İ.A., Malszycki, M., Lübke, A.K., Schade, C., Meierhofer, D., and Aktaş, T. (2020). SON and SRRM2 are essential for nuclear speckle formation. Elife 9, e60579.

28. Wen, L., Arias, M.A., Fan, X., Xie, J., Paul, S., Liao, S.E., Sobczyk, M., Regev, O., and Fei, J. (2026). RNA localization to nuclear speckles follows splicing logic. Nucleic Acids Res. 54, gkag174.

29. Alexander, K.A., Yu, R., Skuli, N., Coffey, N.J., Nguyen, S., Faunce, C.L., Huang, H., Dardani, I.P., Good, A.L., Lim, J., et al. (2025). Nuclear speckles regulate functional programs in cancer. Nat. Cell Biol. 27, 322–335.

30. Chaturvedi, P., Ghosh, P., Zhang, L., Zhang, M., Zhao, H., and Belmont, A.S. (2026). Acidic transcription factors position the genome at nuclear speckles through transcription-dependent and -independent mechanisms. Mol. Cell 86, 3431–3448.e9.

31. Yu, R., Roseman, S., Siegenfeld, A.P., Gardner, Z., Nguyen, S.C., Tran, K.A., Joyce, E.F., Jain, R., Liau, B.B., Krantz, I.D., et al. (2025). CTCF/RAD21 organize the ground state of chromatin-nuclear speckle association. Nat. Struct. Mol. Biol. 32, 1069–1080.

32. Quinodoz, S.A., Ollikainen, N., Tabak, B., Palla, A., Schmidt, J.M., Detmar, E., Lai, M.M., Shishkin, A.A., Bhat, P., Takei, Y., et al. (2018). Higher-order inter-chromosomal hubs shape 3D genome organization in the nucleus. Cell 174, 744–757.e24.

33. Palikyras, S., Varamogiani-Mamatsi, V., Zhu, Y., Mizi, A., Ramasamy, S., Grini, J.V., Liebermann, I., Stavropoulou, A., Puig, A.P., Ogurlu-Tank, T.S., et al. (2026). Senescent cells cluster CTCF on nuclear speckles to instruct an alternative splicing program. Nat. Aging 6, 1592–1610.

34. Klonizakis, A., Alcoverro-Bertran, M., Massó, P., Thomas, J., de Andrés-Aguayo, L., Wei, X., Varamogianni-Mamatsi, V., Nikolaou, C., and Graf, T. (2025). Synergistic and antagonistic activities of IRF8 and FOS enhancer pairs during an immune-cell fate switch. EMBO J. 44, 2025–2055.

35. Greenberg, M.E., and Ziff, E.B. (1984). Stimulation of 3T3 cells induces transcription of the c-fos proto-oncogene. Nature 311, 433–438.

36. Liu, Z., Merkurjev, D., Yang, F., Li, W., Oh, S., Friedman, M.J., Song, X., Zhang, F., Ma, Q., Ohgi, K.A., et al. (2014). Enhancer activation requires trans-recruitment of a mega transcription factor complex. Cell 159, 358–373.

37. Hah, N., Danko, C.G., Core, L., Waterfall, J.J., Siepel, A., Lis, J.T., and Kraus, W.L. (2011). A rapid, extensive, and transient transcriptional response to estrogen signaling in breast cancer cells. Cell 145, 622–634.

38. Chen, L.-F., Lin, Y.T., Gallegos, D.A., Hazlett, M.F., Gómez-Schiavon, M., Yang, M.G., Kalmeta, B., Zhou, A.S., Holtzman, L., Gersbach, C.A., et al. (2019). Enhancer histone acetylation modulates transcriptional bursting dynamics of neuronal activity-inducible genes. Cell Rep. 26, 1174–1188.e5.

39. Joo, J.-Y., Schaukowitch, K., Farbiak, L., Kilaru, G., and Kim, T.-K. (2016). Stimulus-specific combinatorial functionality of neuronal c-fos enhancers. Nat. Neurosci. 19, 75–83.

40. Hezroni, H., Koppstein, D., Schwartz, M.G., Avrutin, A., Bartel, D.P., and Ulitsky, I. (2015). Principles of long noncoding RNA evolution derived from direct comparison of transcriptomes in 17 species. Cell Rep. 11, 1110–1122.

41. Dalton, S., and Treisman, R. (1992). Characterization of SAP-1, a protein recruited by serum response factor to the c-fos serum response element. Cell 68, 597–612.

42. Shaw, P.E., Schröter, H., and Nordheim, A. (1989). The ability of a ternary complex to form over the serum response element correlates with serum inducibility of the human c-fos promoter. Cell 56, 563–572.

43. Schmidt, D., Schwalie, P.C., Wilson, M.D., Ballester, B., Gonçalves, A., Kutter, C., Brown, G.D., Marshall, A., Flicek, P., and Odom, D.T. (2012). Waves of retrotransposon expansion remodel genome organization and CTCF binding in multiple mammalian lineages. Cell 148, 335–348.

44. Hah, N., Murakami, S., Nagari, A., Danko, C.G., and Kraus, W.L. (2013). Enhancer transcripts mark active estrogen receptor binding sites. Genome Res. 23, 1210–1223.

45. Gil, N., and Ulitsky, I. (2018). Production of Spliced Long Noncoding RNAs Specifies Regions with Increased Enhancer Activity. Cell Systems 7, 537–547.e3.

46. Lee, J.-S., and Mendell, J.T. (2020). Antisense-mediated transcript knockdown triggers premature transcription termination. Mol. Cell 77, 1044–1054.e3.

47. Díaz, N., Kruse, K., Erdmann, T., Staiger, A.M., Ott, G., Lenz, G., and Vaquerizas, J.M. (2018). Chromatin conformation analysis of primary patient tissue using a low input Hi-C method. Nat. Commun. 9, 4938.

48. Bak, J.H., Kim, M.H., Liu, L., and Hyeon, C. (2021). A unified framework for inferring the multi-scale organization of chromatin domains from Hi-C. PLoS Comput. Biol. 17, e1008834.

49. Robinson, M.D., and Oshlack, A. (2010). A scaling normalization method for differential expression analysis of RNA-seq data. Genome Biol. 11, R25.

50. Chen, Y., Chen, L., Lun, A.T.L., Baldoni, P.L., and Smyth, G.K. (2025). edgeR v4: powerful differential analysis of sequencing data with expanded functionality and improved support for small counts and larger datasets. Nucleic Acids Res. 53, gkaf018.

51. Lun, A.T.L., and Smyth, G.K. (2015). diffHic: a Bioconductor package to detect differential genomic interactions in Hi-C data. BMC Bioinformatics 16, 258.

52. Shanel, G., Hsieh, T.-H.S., Cattoglio, C., Haham, H.A., Chou, H.-J., Li, J.Z., Shamir, R., Darzacq, X., and Elkon, R. (2025). Transcriptional programs of cell identity and p53-induced stress responses are associated with distinctive features of spatial genome organization. Nucleic Acids Res. 53, gkaf607.

53. Chan, B., and Rubinstein, M. (2024). Activity-driven chromatin organization during interphase: Compaction, segregation, and entanglement suppression. Proc. Natl. Acad. Sci. U. S. A. 121, e2401494121.

54. Goel, V.Y., Aboreden, N.G., Jusuf, J.M., Zhang, H., Mori, L.P., Mirny, L.A., Blobel, G.A., Banigan, E.J., and Hansen, A.S. (2025). Dynamics of microcompartment formation at the mitosis-to-G1 transition. Nat. Struct. Mol. Biol. 32, 2614–2627.

55. Salari, H., Fourel, G., and Jost, D. (2024). Transcription regulates the spatio-temporal dynamics of genes through micro-compartmentalization. Nat. Commun. 15, 5393.

56. Hsieh, T.-H.S., Weiner, A., Lajoie, B., Dekker, J., Friedman, N., and Rando, O.J. (2015). Mapping nucleosome resolution chromosome folding in yeast by Micro-C. Cell 162, 108–119.

57. Hsieh, T.-H.S., Cattoglio, C., Slobodyanyuk, E., Hansen, A.S., Rando, O.J., Tjian, R., and Darzacq, X. (2020). Resolving the 3D landscape of transcription-linked mammalian chromatin folding. Mol. Cell 78, 539–553.e8.

58. Chen, Y., Zhang, Y., Wang, Y., Zhang, L., Brinkman, E.K., Adam, S.A., Goldman, R., van Steensel, B., Ma, J., and Belmont, A.S. (2018). Mapping 3D genome organization relative to nuclear compartments using TSA-Seq as a cytological ruler. J. Cell Biol. 217, 4025–4048.

59. Biayna, J., Baranovskii, A., Chaudhuri, A., Paladin, M., Erdem, B., Keller, L.-E., Barutcu, A.R., Dimmeler, S., Marsico, A., and Dumbović, G. (2026). The periphery of nuclear speckles defines a spatially and temporally regulated compartment of long-lived intron-retained RNAs that resolves during mitosis. Nat. Cell Biol. 28, 1857–1874.

60. Konermann, S., Brigham, M.D., Trevino, A.E., Joung, J., Abudayyeh, O.O., Barcena, C., Hsu, P.D., Habib, N., Gootenberg, J.S., Nishimasu, H., et al. (2015). Genome-scale transcriptional activation by an engineered CRISPR-Cas9 complex. Nature. 10.1038/nature14136.

61. Perry, R.B.-T., Hezroni, H., Goldrich, M.J., and Ulitsky, I. (2018). Regulation of Neuroregeneration by Long Noncoding RNAs. Mol. Cell 72, 553–567.e5.

62. Ben-Tov Perry, R., Tsoory, M., Tolmasov, M., and Ulitsky, I. (2023). Silc1 long noncoding RNA is an immediate-early gene promoting efficient memory formation. Cell Rep. 42, 113168.

63. Isoda, T., Moore, A.J., He, Z., Chandra, V., Aida, M., Denholtz, M., Piet van Hamburg, J., Fisch, K.M., Chang, A.N., Fahl, S.P., et al. (2017). Non-coding transcription instructs chromatin folding and compartmentalization to dictate enhancer-promoter communication and T cell fate. Cell 171, 103–119.e18.

64. Krawczyk, M., and Emerson, B.M. (2014). p50-associated COX-2 extragenic RNA (PACER) activates COX-2 gene expression by occluding repressive NF-kappaB complexes. Elife 3, e01776.

65. Arner, E., Daub, C.O., Vitting-Seerup, K., Andersson, R., Lilje, B., Drabløs, F., Lennartsson, A., Rönnerblad, M., Hrydziuszko, O., Vitezic, M., et al. (2015). Transcribed enhancers lead waves of coordinated transcription in transitioning mammalian cells. Science 347, 1010–1014.

66. Heinz, S., Texari, L., Hayes, M.G.B., Urbanowski, M., Chang, M.W., Givarkes, N., Rialdi, A., White, K.M., Albrecht, R.A., Pache, L., et al. (2018). Transcription elongation can affect genome 3D structure. Cell 174, 1522–1536.e22.

67. Hu, S., Lv, P., Yan, Z., and Wen, B. (2019). Disruption of nuclear speckles reduces chromatin interactions in active compartments. Epigenetics Chromatin 12, 43.

68. Roseman, S.A., Siegenfeld, A.P., Lee, C., Lue, N.Z., Waterbury, A.L., and Liau, B.B. (2025). DNA methylation insulates genic regions from CTCF loops near nuclear speckles. Elife 13, RP102930.

69. Alexander, K.A., Coté, A., Nguyen, S.C., Zhang, L., Gholamalamdari, O., Agudelo-Garcia, P., Lin-Shiao, E., Tanim, K.M.A., Lim, J., Biddle, N., et al. (2021). p53 mediates target gene association with nuclear speckles for amplified RNA expression. Mol. Cell 81, 1666–1681.e6.

70. Sung, H.-M., Schott, J., Boss, P., Lehmann, J.A., Hardt, M.R., Lindner, D., Messens, J., Bogeski, I., Ohler, U., and Stoecklin, G. (2023). Stress-induced nuclear speckle reorganization is linked to activation of immediate early gene splicing. J. Cell Biol. 222, e202111151.

71. Choi, H.M.T., Schwarzkopf, M., Fornace, M.E., Acharya, A., Artavanis, G., Stegmaier, J., Cunha, A., and Pierce, N.A. (2018). Third-generation in situ hybridization chain reaction: multiplexed, quantitative, sensitive, versatile, robust. Development 145, dev165753.

72. Potapova, T.A., Unruh, J.R., Conkright-Fincham, J., Banks, C.A.S., Florens, L., Schneider, D.A., and Gerton, J.L. (2023). Distinct states of nucleolar stress induced by anticancer drugs. Elife 12, RP88799.

73. Ramasamy, S., Zhu, Y., Karpinska, M., Flyamer, I., Söding, J., and Oudelaar, M. (2026). Analysis of multi-way chromatin interactions at sub-nucleosome resolution identifies synergistic nano-scale interactions within super-enhancers. bioRxiv. 10.64898/2026.05.21.726779.

74. Hamley, J.C., Li, H., Denny, N., Downes, D., and Davies, J.O.J. (2023). Determining chromatin architecture with Micro Capture-C. Nat. Protoc. 18, 1687–1711.

