## Supplementary material for "The long noncoding RNA *FOSIL* shapes the dynamics of *FOS* induction": FOSIL_Supplementary_Methods

#### Key resources table

| Reagent or resource | Source | Identifier or status |
|---|---|---|
| MCF7 cells | Ulitsky laboratory | MCF7 laboratory stock |
| LNA-GapmeRs | Qiagen | Qiagen Antisense LNA-GapmeRs; CTRL: Negative Control B (LG00000001); L1 and L2 target FOSIL |
| CRISPRa guides / constructs | This study | SAM: dCas9-VP64/MS2-p65-HSF1; empty guide vector; *FOSIL* promoter sgRNAs (sequences in main Methods) |
| Ectopic *FOSIL* construct | This study | pcDNA3.1(+), CMV promoter; Twist-synthesized four-exon FOSIL insert |
| *FOSIL* intronic HCR probes | Molecular Instruments | HSS-LINC01220introns-B4 |
| *FOS* intron/exon and *MALAT1* HCR probes | Molecular Instruments | FOS introns-B2, FOS exons-B5 and MALAT1-B2; combinations described in main Methods |
| HCR amplifiers | Molecular Instruments | L/M series: B2-Alexa Fluor 488; B4-Alexa Fluor 647; B5-Alexa Fluor 546 |
| *FOS* DNA FISH probe | Empire Genomics | FOS-20-OR; orange |
| Mouse anti-SC35 | Abcam | SC-35; Abcam ab11826 |
| Goat anti-mouse Alexa 488 | Abcam | ab150113 |
| Digitonin | Sigma-Aldrich | D141-100MG |
| Concanavalin A beads | EpiCypher | 21-1411 |
| Micrococcal nuclease | New England Biolabs | M0247 |
| DNeasy Blood & Tissue kit | QIAGEN | 69506 |
| NEBNext Ultra II DNA kit | New England Biolabs | E7645L |
| Custom 70-mer capture panels | Twist Bioscience | TE-95053285 |
| ProLong Diamond | Invitrogen | ProLong Diamond Antifade Mountant |

#### Capture Micro-C processing and contact analysis

Paired-end reads were aligned to GRCh38 (primary assembly) with BWA-MEM 0.7.19 (-SP) [75], parsed, sorted and deduplicated with pairtools 1.1.3 (parse2 --min-mapq 30; dedup --max-mismatch 1) [21,76], and binned into cooler 0.10.4 matrices at 250 bp, 500 bp and 1 kb over the captured window (chr14:75,048,709-75,388,707) [77]. Replicate captures of the same WT library were merged prior to deduplication. For contact calling, replicates were summed within each condition and state. The maps were balanced by iterative correction in cooler [77,78], using cis reads and excluding the first two diagonals and low-coverage bins. Mustache 1.3.3 was run at 1 kb on each state map and at 1 kb and 250 bp on the map pooled across all 48 CTRL, L1 and L2 libraries (sparsity threshold 0.7, P-value threshold 0.1) [79]. Contacts detected across all four states in CTRL were merged across adjacent pixels and ranked by serum contact strength, defined as the balanced count in a 3 × 3-kb box centered on the contact divided by the distance-matched expected count [54,80]. Comparison with public genome-wide MCF7 Micro-C (GSE278978) [52] used unbalanced 2-kb matrices. Contact hubs were defined in 5-kb bins as positions with contact-end counts at least 1.5 SD above the window mean [81], assessed against 10,000 random length-preserving placements.

Insulation scores were computed on unbalanced 1-kb matrices following Crane et al. [82] as the log2 ratio of the mean read pairs in the 10 × 10-kb square flanking each 1-kb bin (excluding the bin itself) to the library's median score across the captured window [80]. Domain boundaries were defined at local insulation minima reaching at least the 93rd percentile of topographic prominence across CTRL libraries (chr14:75,259,500 for |e2 core, 75,277,500 for e4|*FOS*, 75,283,500 for *FOS*|*FOSIL*, 75,294,500 for |*FOSIL*$_{TSS}$, and 75,307,500 for |CTCF$_R$). Interval scores were calculated as the mean insulation score across interior 1-kb bins of the *FOS* gene (75,277.5-75,283.5 kb) and the intergenic *FOS→FOSIL* interval (75,283.5-75,294.5 kb).

Differential interactions and insulation changes were tested on deduplicated read-pair counts in 3 × 3-kb contact boxes, 10 × 10-kb insulation squares and interior interval bins using diffHic 1.42.0 [51]. Libraries were normalized by TMM [49] on bin pairs with mean count ≥ 5 and evaluated with edgeR 4.8.2 quasi-likelihood F-tests using robust dispersion estimation (~ line * state + capture) [50], with significance called at Benjamini-Hochberg FDR < 1%. Significantly changing *FOS*-promoter contacts and insulation positions were grouped by temporal profile using k-means clustering (k = 4 for contacts, k = 3 for insulation; 1,000 random starts) on standardized four-state differences.

#### Genomic annotation and public datasets

All coordinates refer to GRCh38 (*FOS*-201 MANE Select; *FOSIL* NR_038421.1 RefSeq). MCF7 chromatin and transcription-factor profiles and candidate cis-regulatory elements (ENCSR891IDS) were obtained from ENCODE [83]. FANTOM5 hg38 CAGE tags [5] and estradiol-treated MCF7 GRO-seq datasets (GSE27463 and GSE60272, lifted from hg18 to GRCh38 with UCSC liftOver) [36,37] were binned at 50 bp and normalized to library depth. Plus-strand read-through was quantified over chr14:75,285,191-75,293,903 (beginning 2 kb downstream of *FOS*) relative to the *FOS* gene body (chr14:75,278,828-75,282,230). *FOSIL* splice-junction usage in GTEx v8, TCGA and mouse SRA (srav1m) RNA-seq compilations was queried via Snaptron and recount3 [84,85].

#### Evolutionary sequence conservation and motif analysis

Synteny and same-strand noncoding candidates in the *TMED10-FOS-JDP2* interval (within 60 kb downstream of *FOS*) were surveyed across 15 vertebrates using current NCBI RefSeq and Ensembl annotations and UCSC CpG islands. Rodent orthologs at the aligned *FOSIL* exon-3 position were obtained from Ensembl release 116 (*Gm60138*, GRCm39) and RefSeq (*LOC148005395*, GRCr8), and mammalian CTCF occupancy was evaluated across 42 ENCODE mouse ChIP-seq datasets [83] and five-species liver ChIP-seq (ArrayExpress E-MTAB-437) [43]. Base-level conservation and conserved elements were obtained from UCSC phyloP and phastCons tracks (phyloP100way, phyloP470wayBW, phyloP30way, phyloP447wayPrimates) [86,87] and compared against 10,000 random length-matched intervals within chr14:75,230,001-75,330,000 (Benjamini-Hochberg adjusted). Across 18,534 GENCODE v42 basic lncRNAs (≥ 200-bp exon union with ≥ 90% scored bases), genes were ranked by mean 100-way vertebrate phyloP, and sequence-ortholog counts across 20 species were determined as described in Hezroni et al. [40].

Transcription-factor motifs were scanned with JASPAR 2024 matrices [88] for CTCF (MA0139.1, relative score ≥ 0.80), SRF (MA0083.3) and ELK1 (MA0028.3, within 40 bp of exact CC(A/T)6GG CArG boxes), and intact elements were scored across the UCSC 470-way mammalian and 100-way vertebrate alignments. MCF7 ChIP-seq peaks overlapping the *FOSIL* exon-3 CTCF site and the *FOSIL* and *FOS* SREs were ranked genome-wide and within 500 kb across eight ENCODE CTCF, two RAD21, and the MCF7 SRF and ELK1 conservative IDR peak sets.

#### RNA-seq alignment, quantification and specificity analysis

Paired-end RNA-seq reads (2 × 61 bp) were aligned to GRCh38 with STAR [89]. Gene and *FOSIL* counts (across the 2,007-bp NR_038421.1 span and 490-bp exon union) were quantified with STAR and featureCounts 2.0.6 [90] against GENCODE v42 and MANE v1.0 annotations and normalized by DESeq2 median-of-ratios size factors [91].

Differential gene expression against time-matched CTRL was tested with DESeq2 Wald tests (PyDESeq2 0.5.4) [91,92] with Benjamini-Hochberg adjustment, and transcriptome-wide specificity was evaluated against a $|\log2FC| > 1$ threshold (adjusted $P < 0.05$). *FOSIL* coverage reduction across the five time points was evaluated with Gart's one-sided exact conditional test (mid-P) [93]. Candidate GapmeR off-targets were identified by full-length sense-strand alignment to GRCh38 MANE v1.0 genes with ≤ 3 mismatches, and essentiality across 1,178 cell lines (53 breast lines) was obtained from DepMap 24Q4 CRISPR Chronos scores (dependency probability > 0.5) [94,95].

#### Quantification and statistical analysis

##### RT-qPCR and flow cytometry

RT-qPCR ΔCt values across the GapmeR time course were analyzed by two-way ANOVA (condition × time) followed by Dunnett's test against time-matched CTRL; serum baseline (S) was analyzed by Dunnett's test and the CRISPRa time course by pooled-variance t-tests against time-matched EMPTY, and ectopic expression by Welch's t-tests with Holm correction. Flow cytometry data (9,159-23,823 cells per pooled sample) were evaluated by one-sided minimum-effect tests against a standardized threshold of 0.2 (Cohen's d on log10 FOS intensity; Cohen's h on arcsine-square-root-transformed positive fractions) with Holm correction.

##### RNA imaging and spatial measurements

Nuclei were segmented from DAPI maximum-intensity projections with Cellpose [96], RNA-HCR transcription active sites (TAS) were detected with StarDist [97] and custom Fiji/ImageJ macros (with blinded manual curation of a subset of images), and DNA-FISH loci were detected with Spotiflow [98]. In the M series, *MALAT1* speckle masks were defined on average-intensity projections at 2× the mean nuclear *MALAT1* intensity. Across both imaging series, 0.54 µm rolling-ball background subtraction was applied before thresholding. Each DNA-FISH locus-to-speckle distance was divided by the mean distance from nuclear pixels to the nearest SC35 speckle mask in the same nucleus.

In untransfected WT cells (L series), sites in nuclei with at least two TAS were grouped by preFOSIL intensity relative to the mean at each time point, calculated across sites or within each nucleus. Log10 fluorescence was compared between groups using a linear mixed model (lme4) with replicate as a fixed effect, random intercepts for image and nucleus, and a random slope for the high/low contrast by image. P values from likelihood-ratio tests were adjusted by the Holm method. Comparisons of per-site fluorescence, active-site counts per nucleus, and locus-to-speckle distances between GapmeR or CRISPRa conditions and matched controls (or between top and bottom preFOSIL quarters) used two-sided Wilcoxon rank-sum tests; full distance distributions were compared by two-sample Kolmogorov-Smirnov tests, and speckle-proximal fractions (≤ 0.15 µm and ≤ 0.32 µm) by two-sided Fisher's exact tests.

##### Statistical reporting

Biological replicates were independently grown or transfected cultures. Capture Micro-C included four replicates per GapmeR condition and two WT replicates at each time point (48 GapmeR and 8 WT libraries). RT-qPCR used three replicates, or two for the CRISPRa time course. RNA-seq, each imaging series and the proliferation assay used two independent experiments. For flow cytometry, three independently transfected cultures were pooled per condition and time point. Sample sizes, summary metrics and P-value thresholds are reported in the figure legends.

### Supplementary References

References 1–74 are listed in the main manuscript.

75. Li, H. (2013). Aligning sequence reads, clone sequences and assembly contigs with BWA-MEM. arXiv [q-bio.GN]. https://doi.org/10.48550/arXiv.1303.3997.

76. Open2C, Abdennur, N., Fudenberg, G., Flyamer, I.M., Galitsyna, A.A., Goloborodko, A., Imakaev, M., and Venev, S.V. (2024). Pairtools: From sequencing data to chromosome contacts. PLoS Comput. Biol. *20*, e1012164.

77. Abdennur, N., and Mirny, L.A. (2020). Cooler: scalable storage for Hi-C data and other genomically labeled arrays. Bioinformatics *36*, 311–316.

78. Imakaev, M., Fudenberg, G., McCord, R.P., Naumova, N., Goloborodko, A., Lajoie, B.R., Dekker, J., and Mirny, L.A. (2012). Iterative correction of Hi-C data reveals hallmarks of chromosome organization. Nat. Methods *9*, 999–1003.

79. Roayaei Ardakany, A., Gezer, H.T., Lonardi, S., and Ay, F. (2020). Mustache: multi-scale detection of chromatin loops from Hi-C and Micro-C maps using scale-space representation. Genome Biol. *21*, 256.

80. Open2C, Abdennur, N., Abraham, S., Fudenberg, G., Flyamer, I.M., Galitsyna, A.A., Goloborodko, A., Imakaev, M., Oksuz, B.A., Venev, S.V., et al. (2024). Cooltools: Enabling high-resolution Hi-C analysis in Python. PLoS Comput. Biol. *20*, e1012067.

81. Huang, J., Li, K., Cai, W., Liu, X., Zhang, Y., Orkin, S.H., Xu, J., and Yuan, G.-C. (2018). Dissecting super-enhancer hierarchy based on chromatin interactions. Nat. Commun. *9*, 943.

82. Crane, E., Bian, Q., McCord, R.P., Lajoie, B.R., Wheeler, B.S., Ralston, E.J., Uzawa, S., Dekker, J., and Meyer, B.J. (2015). Condensin-driven remodelling of X chromosome topology during dosage compensation. Nature *523*, 240–244.

83. ENCODE Project Consortium, Moore, J.E., Purcaro, M.J., Pratt, H.E., Epstein, C.B., Shoresh, N., Adrian, J., Kawli, T., Davis, C.A., Dobin, A., et al. (2020). Expanded encyclopaedias of DNA elements in the human and mouse genomes. Nature *583*, 699–710.

84. Wilks, C., Gaddipati, P., Nellore, A., and Langmead, B. (2018). Snaptron: querying splicing patterns across tens of thousands of RNA-seq samples. Bioinformatics *34*, 114–116.

85. Wilks, C., Zheng, S.C., Chen, F.Y., Charles, R., Solomon, B., Ling, J.P., Imada, E.L., Zhang, D., Joseph, L., Leek, J.T., et al. (2021). recount3: summaries and queries for large-scale RNA-seq expression and splicing. Genome Biol. *22*, 323.

86. Siepel, A., Bejerano, G., Pedersen, J.S., Hinrichs, A.S., Hou, M., Rosenbloom, K., Clawson, H., Spieth, J., Hillier, L.W., Richards, S., et al. (2005). Evolutionarily conserved elements in vertebrate, insect, worm, and yeast genomes. Genome Res. *15*, 1034–1050.

87. Pollard, K.S., Hubisz, M.J., Rosenbloom, K.R., and Siepel, A. (2010). Detection of nonneutral substitution rates on mammalian phylogenies. Genome Res. *20*, 110–121.

88. Rauluseviciute, I., Riudavets-Puig, R., Blanc-Mathieu, R., Castro-Mondragon, J.A., Ferenc, K., Kumar, V., Lemma, R.B., Lucas, J., Chèneby, J., Baranasic, D., et al. (2024). JASPAR 2024: 20th anniversary of the open-access database of transcription factor binding profiles. Nucleic Acids Res. *52*, D174–D182.

89. Dobin, A., Davis, C.A., Schlesinger, F., Drenkow, J., Zaleski, C., Jha, S., Batut, P., Chaisson, M., and Gingeras, T.R. (2013). STAR: ultrafast universal RNA-seq aligner. Bioinformatics *29*, 15–21.

90. Liao, Y., Smyth, G.K., and Shi, W. (2014). featureCounts: an efficient general purpose program for assigning sequence reads to genomic features. Bioinformatics *30*, 923–930.

91. Love, M.I., Huber, W., and Anders, S. (2014). Moderated estimation of fold change and dispersion for RNA-seq data with DESeq2. Genome Biol. *15*, 550.

92. Muzellec, B., Teleńczuk, M., Cabeli, V., and Andreux, M. (2023). PyDESeq2: a python package for bulk RNA-seq differential expression analysis. Bioinformatics *39*, btad547.

93. Gart, J.J. (1978). The analysis of ratios and cross-product ratios of poisson variates with application to incidence rates. Commun. Stat. Theory Methods *7*, 917–937.

94. Broad DepMap (2024). DepMap 24Q4 Public. (Figshare+). https://doi.org/10.25452/FIGSHARE.PLUS.27993248.V1.

95. Dempster, J.M., Boyle, I., Vazquez, F., Root, D.E., Boehm, J.S., Hahn, W.C., Tsherniak, A., and McFarland, J.M. (2021). Chronos: a cell population dynamics model of CRISPR experiments that improves inference of gene fitness effects. Genome Biol. *22*, 343.

96. Stringer, C., Wang, T., Michaelos, M., and Pachitariu, M. (2021). Cellpose: a generalist algorithm for cellular segmentation. Nat. Methods *18*, 100–106.

97. Schmidt, U., Weigert, M., Broaddus, C., and Myers, G. (2018). Cell detection with star-convex polygons. In Medical Image Computing and Computer Assisted Intervention – MICCAI 2018 Lecture Notes in Computer Science. (Springer International Publishing), pp. 265–273.

98. Dominguez Mantes, A., Herrera, A., Khven, I., Schlaeppi, A., Kyriacou, E., Tsissios, G., Skoufa, E., Santangeli, L., Buglakova, E., Durmus, E.B., et al. (2025). Spotiflow: accurate and efficient spot detection for fluorescence microscopy with deep stereographic flow regression. Nat. Methods *22*, 1495–1504.
